# Rapid Orthographic and Delayed Phonological Processing: ERP and Oscillatory Evidence from Masked Priming in Korean

**DOI:** 10.64898/2026.03.05.709970

**Authors:** Joonwoo Kim, Solbin Lee, Kichun Nam

**Author notes:** Corresponding author. E-mail address (K. Nam).

## Abstract

A central question in visual word recognition concerns whether orthographic and phonological codes are coordinated sequentially or in parallel during lexical access. Korean Hangul, an alpha-syllabic writing system with morphophonemic spelling principles, allows independent manipulation of orthographic and phonological syllable overlap. In a masked priming lexical decision task with EEG (*N* = 30), we contrasted orthographically identical primes (e.g., 식-식량), phonologically overlapping primes (e.g., 싱-식량), and unrelated primes. Event-related potentials and time-frequency representations (theta: 4–8 Hz, lower beta: 13–20 Hz, upper beta: 20–30 Hz) were analyzed to capture both evoked and oscillatory neural dynamics. Orthographic priming produced a cascade of facilitative effects: early fronto-central P200 enhancement (150–250 ms) with upper beta synchronization (30–290 ms), followed by centro-parietal N400 reduction (350–550 ms) with frontal theta suppression (400–730 ms), and behavioral facilitation. Phonological priming, by contrast, elicited an inhibitory behavioral effect and sustained lower beta activity over central regions (310–590 ms) but produced no early electrophysiological modulation, consistent with lexical competition. This spatiotemporal dissociation suggests that orthographic syllable processing can emerge at early stages and cascades into later lexical-level processing, whereas phonological syllable effects are confined to later processing stages. These findings provide support for a sequential or cascaded account of orthographic-phonological coordination, as predicted by dual-route models, while challenging strong forms of parallel activation, and suggest that the alpha-syllabic structure of Korean may enable a processing strategy in which orthographic parsing serves as an efficient entry route to the lexicon.

## 1. Introduction

Understanding how orthographic and phonological codes are coordinated during lexical access has been a long-standing question in visual word recognition research. Dual-route models such as the Dual Route Cascaded (DRC) model posit distinct processing pathways—a direct orthographic route that maps visual input onto whole-word representations and an indirect phonological route that assembles phonological codes prior to lexical access (Coltheart et al., 2001; Perry et al., 2007)—predicting that orthographic analysis precedes phonological assembly. Interactive activation frameworks such as the Bi-Modal Interactive Activation Model (BIAM; Diependaele et al., 2010; Grainger & Holcomb, 2009), by contrast, propose that orthographic and phonological codes are activated rapidly and in parallel, building on the foundational Interactive Activation Model (IAM; McClelland & Rumelhart, 1981). Sublexical units occupy a central role in this debate, as they mediate between orthographic input and lexical access and represent the earliest processing level at which the two codes interact. Two related questions have motivated much of this research: whether sublexical effects reflect orthographic or phonological representations, and whether they originate at early form-encoding stages or at later lexical stages during competitive activation. However, since alphabetic writing systems inherently confound orthographic and phonological syllable structure, independent manipulation of the two has remained elusive, leaving both questions yet to be resolved.

### 1.1. Orthographic and Phonological Sublexical Variables

To examine the respective contributions of orthographic and phonological information, a range of sublexical variables have been investigated in visual word recognition. Orthographic variables, such as bigram frequency, orthographic neighborhood density, and letter position encoding, have consistently been reported to produce facilitatory effects on lexical access, suggesting that frequent or familiar orthographic patterns reduce the processing demands of visual form analysis (Andrews, 1997; Grainger et al., 2005). Phonological variables, by contrast, tend to produce inhibitory effects: phonological neighborhood density slows lexical decisions as the number of phonologically similar words increases, consistent with lexical competition driven by phonological activation (Grainger et al., 2005; Luce & Pisoni, 1998).

The representational distinction intersects with a longstanding debate about the role of phonological codes in visual word recognition. According to dual-route and cascaded architectures (e.g., Coltheart et al., 2001), orthographic form analysis provides a fast, direct pathway to the lexicon, with phonological assembly proceeding through a slower indirect route. An alternative class of accounts maintains that phonological codes are computed rapidly and obligatorily, serving a functional role in lexical access regardless of task or orthographic system (Brysbaert, 2001; Lukatela et al., 1998, 2001). Evidence from masked priming studies has been equivocal: early demonstrations of phonological effects in French (Ferrand & Grainger, 1992, 1993) were not consistently replicated in deeper orthographies such as English (see Rastle & Brysbaert, 2006, for review). However, subsequent work has shown that null results reflected inadequate control of orthographic neighborhood competition rather than a genuine absence of early phonological activation (Davis & Lupker, 2006; Rastle & Brysbaert, 2006). The debate has accordingly shifted from whether phonological codes are activated early to the conditions under which such activation becomes detectable, a question that remains open.

Within this broader debate, the syllable is of particular interest as a sublexical unit that straddles both orthographic and phonological dimensions. Syllable frequency effects (SFEs), in which words beginning with high-frequency syllables are recognized more slowly than those with low-frequency syllables, have been documented across multiple alphabetic languages, including Spanish (Carreiras et al., 1993; Perea & Carreiras, 1998), German (Conrad & Jacobs, 2004), and French (Mathey & Zagar, 2002). These inhibitory SFEs are consistent with the phonological pattern described above, that high-frequency syllables activate a larger set of lexical competitors, slowing target recognition via lateral inhibition (Grainger & Jacobs, 1996; McClelland & Rumelhart, 1981). Computational models such as the MROM-S (Conrad et al., 2010) and CDP++ (Perry et al., 2010) have formalized this mechanism by incorporating a syllabic parser that bridges sublexical form analysis and lexical access along the phonological route. Taken together, the broad pattern across sublexical variables suggests an asymmetry in which orthographic information facilitates processing via rapid form-level activation, while phonological information, including syllabic units, tends to generate competition and inhibition.

### 1.2. Korean Hangul: Dissociating Orthographic and Phonological Syllables

Korean Hangul provides a unique opportunity to address both of these questions—representational basis and processing stage—within a single experimental paradigm. Unlike alphabetic scripts, Hangul is an alpha-syllabic writing system in which individual phonemes are spatially arranged into compact syllabic blocks that function as orthographic units (Pae, 2011; Simpson & Kang, 2004). The syllable 곰 /kom/, for instance, comprises three graphemes (ㄱ-ㅗ-ㅁ) arranged in a two-dimensional configuration, forming a single visual unit. This spatial arrangement creates strong perceptual grouping and precise orthographic encoding, as evidenced by the absence of transposed-letter effects that are commonly found in alphabetic scripts (C. H. Lee & Taft, 2009).

Crucially, Korean orthography follows morphophonemic principles that preserve morphological boundaries in spelling even when pronunciation changes occur across syllable boundaries. This property allows systematic dissociation of orthographic from phonological syllable overlap within the same word. For example, the prime 식 / ik / and target 식량 / iŋ aŋ/ share the same orthographic syllable (식), but their phonological representations differ due to resyllabification (/ ik / vs. / iŋ/). Conversely, the prime 싱 / iŋ/ and target 식량 / iŋ aŋ/ share phonological overlap (/ iŋ/) but are orthographically distinct (싱 vs. 식). This crossed design provides the experimental leverage to independently manipulate orthographic and phonological syllable relatedness—a constraint that is difficult to achieve systematically in alphabetic orthographies. Although some alphabetic scripts permit partial dissociation of orthographic and phonological syllables (e.g., Álvarez et al., 2004), Korean morphophonemic spelling provides a more extensive and systematic basis for fully crossed designs—while simultaneously enabling examination of whether effects arise at early or later processing stages.

These properties position Korean as an ideal test of the conditions identified in the reframed phonological recoding debate. Hangul’s morphophonemic orthography and compact block structure create a processing environment in which orthographic units are visually salient and immediately available, while phonological codes must be assembled across morphophonemic boundaries. If phonological codes are generated rapidly and automatically, as predicted by phonological recoding accounts, their effects on syllable processing should emerge early, potentially overlapping with the time course of orthographic effects. If, however, phonological assembly is delayed in this writing system, as predicted by the DRC framework, then orthographic and phonological syllable effects should be temporally dissociated, with phonological effects appearing later in the processing stream.

### 1.3. Previous Korean Syllable Processing Research

A central debate in Korean visual word recognition concerns whether phonological information is automatically activated or whether recognition proceeds primarily through orthographic processing. Orthographic syllables have emerged as robust processing units across paradigms. Park (1996) found no facilitation from pseudohomophone primes relative to orthographic controls, suggesting that phonological information is not automatically activated. Bae and Yi (2010) compared orthographically identical first-syllable primes (숙소-숙녀) with phonologically identical primes (숭배-숙녀) and found facilitation only for orthographic overlap.

Similarly, Tae et al. (2015) manipulated prime duration (60 ms, 150 ms) and found significant priming effects only for orthographically matched primes. This orthographic dominance extended to cross-modal priming with auditory primes (Tae et al., 2017), and a recent meta-analysis further confirmed the dominant role of orthographic syllables across paradigms (Lim et al., 2022).

However, the role of phonological syllables remains contested. Some studies reported inhibitory effects of phonological syllables, consistent with lexical competition driven by syllable frequency (Choi et al., 2015; Y. Kwon, 2012; Y. Kwon et al., 2011; Y. Kwon & Lee, 2017). Other studies found null effects (Jin et al., 2018; Y. Kwon & Lee, 2015), and more recent work has reported facilitatory effects, particularly in morphologically complex words, attributed to a fast-guess mechanism (Kim et al., 2023; S. Kwon et al., 2023; S. Lee et al., 2023).

The processing stage of these effects has been directly examined by Choi et al. (2015). Using word primes undergoing phonological assimilation, they replicated the canonical dissociation—phonological overlap yielding inhibition, orthographic overlap yielding facilitation. Critically, nonword primes with identical phonological structure produced null effects in both conditions, suggesting that syllabic competition may be localized to the lexical level in Korean. This finding is consistent with behavioral evidence from alphabetic languages (Carreiras & Perea, 2002; Dominguez et al., 1997) and suggests that the asymmetry between orthographic facilitation and phonological inhibition reflects a difference not only in representational basis but also in processing stage.

Overall, orthographic overlap consistently facilitates recognition across paradigms, whereas phonological overlap yields heterogeneous outcomes shaped by task demands, lexical competition, and morphological structure. Resolving the nature of this asymmetry requires neural measures with sufficient temporal resolution to distinguish the processing stages at which orthographic and phonological syllable effects arise.

### 1.4. ERP Evidence: Processing Stage and Temporal Dynamics

Event-related potentials provide the temporal resolution necessary for investigating whether sublexical effects arise at early form-encoding or later lexical processing stages. The P200 component (150–250 ms), characterized by a fronto-central scalp distribution, has been interpreted as an index of early sublexical form encoding and has been associated with early orthographic–phonological processing, where high-frequency syllables consistently produce reduced P200 amplitudes, interpreted as reflecting more efficient sublexical activation (Barber et al., 2004; Hutzler et al., 2004). The N400 component (300–500 ms), with its characteristic centro-parietal scalp distribution, has been linked to lexical access and semantic integration, with high-frequency syllables producing enhanced negativity attributed to increased lexical competition from a larger neighborhood of words (Carreiras et al., 2005). This P200-facilitation/N400-competition dissociation has been interpreted in terms of a distinction between early sublexical form encoding and later lexical processing. However, several studies have reported SFEs across both time windows simultaneously (Chetail et al., 2012; Goslin et al., 2006), leaving the neural locus unresolved in alphabetic languages.

Korean ERP studies have paralleled these findings while providing additional leverage through the orthographic-phonological dissociation inherent in the writing system. Phonological syllable frequency modulated P200 amplitudes (150–250 ms) in pseudoword recognition, with high-frequency phonological syllables producing greater negativity, suggesting early inhibitory activation (Y. Kwon et al., 2011). The N400 (300–500 ms) was primarily influenced by syllable neighborhood density, with denser neighborhoods producing enhanced negativity reflecting lexical competition (Y. Kwon et al., 2012; Y. Kwon & Lee, 2015). Orthographic syllable frequency, by contrast, produced facilitatory P300 effects (250–350 ms) in morphologically complex nouns (S. Kwon et al., 2024). This dissociation was further supported by masked priming evidence (Y. Lee et al., 2016), where both orthographically and phonologically identical primes produced robust N/P150 effects (100–200 ms; see Grainger & Holcomb, 2009), while phonologically identical primes showed no N250 effects (300–400 ms). Together, Korean ERP evidence suggests an early orthographically driven advantage alongside a delayed phonological competition effect, providing support for a temporal dissociation consistent with distinct processing stages for the two syllable types, though one that warrants further examination with respect to the temporal locus of these effects.

### 1.5. Time-Frequency Analysis: Beyond ERPs

While ERPs reveal the temporal dynamics of word recognition at discrete latency windows, time-frequency analysis exposes oscillatory mechanisms that are not visible in conventional ERP averaging. Distinct frequency bands have been linked to specific cognitive operations: theta oscillations (4–8 Hz) have been associated with lexical retrieval and semantic integration (Bastiaansen et al., 2005, 2008), and beta oscillations (13–30 Hz) have been implicated in top-down prediction and syntactic integration (Lewis & Bastiaansen, 2015; Weiss & Mueller, 2012). Beta oscillations can be further decomposed into lower beta (13–20 Hz), associated with sustained cognitive processing (Kilavik et al., 2013; Pfurtscheller & Lopes da Silva, 1999), and upper beta (20–30 Hz), reflecting faster coordination dynamics during perceptual and representational processing (Engel & Fries, 2010).

Crucially, oscillatory dynamics reflect the coordination and updating of distributed neural representations rather than facilitation per se, providing access to computational processes that are orthogonal to amplitude-based ERP measures (Lewis et al., 2015; Wang et al., 2012). Given that orthographic facilitation and phonological competition are hypothesized to rely on distinct computational mechanisms—rapid perceptual grouping versus sustained assembly and conflict resolution—oscillatory dynamics may offer diagnostic signatures capable of dissociating these processes beyond what ERPs alone can reveal.

### 1.6. The Present Study

The present study employed masked priming with lexical decision and high-density EEG (N = 30) to simultaneously address two questions: (1) whether syllabic priming effects in Korean reflect orthographic or phonological syllable overlap, and (2) whether these effects originate at early form-encoding stages or at later lexical stages. Using the crossed design afforded by Korean morphophonemic spelling, we contrasted orthographically identical primes (e.g., 식-식량 / ik /-/ iŋ aŋ/) with phonologically overlapping primes (e.g., 싱-식량 / iŋ/-/ iŋ aŋ/). Both event-related potentials and time-frequency representations (theta 4–8 Hz, lower beta 13–20 Hz, upper beta 20–30 Hz) were analyzed to capture evoked responses and oscillatory dynamics of syllable priming.

Based on dual-route accounts (Coltheart et al., 2001; Perry et al., 2007) and the alpha-syllabic characteristics of Korean, we derived differential predictions for orthographic and phonological priming across behavioral, electrophysiological, and oscillatory domains. If orthographic syllable overlap facilitates early form-based encoding, it should produce early electrophysiological modulation consistent with enhanced form processing efficiency, reflected in increased P200 amplitudes (150–250 ms) over fronto-central sites, followed by reduced N400 responses (350–550 ms) over centro-parietal sites indexing facilitated lexical–semantic integration. This pattern should be accompanied by facilitative priming effect (i.e., faster latencies relative to unrelated pairs). In contrast, if phonological syllable overlap primarily engages lexical-level assembly and competition processes rather than early visual encoding, early P200 modulation should be absent. Instead, later effects emerging at N400 latencies would be expected, reflecting competitive activation or representational conflict, with an inhibitory priming effect.

Finally, oscillatory dynamics were expected to dissociate the computational mechanisms underlying these effects. Orthographic priming was hypothesized to preferentially modulate upper beta activity (20–30 Hz), consistent with rapid stabilization of perceptual representations. Phonological priming, by contrast, was predicted to recruit sustained theta (4–8 Hz) and lower beta (13–20 Hz) dynamics associated with iterative phonological assembly, lexical competition, and increased coordination of distributed representations.

## 2. Methods

### 2.1. Participants

Participants were 30 healthy, right-handed, and native Korean speakers (17 female; age: 23.33 ± 2.6 years), who had normal or corrected-to-normal vision and did not report any history of psychiatric or neurological disorder. Their handedness was confirmed by Edinburgh Handedness Test scores (Oldfield, 1972; M ± SD = 8.47 ± 1.46).

The sample size of 30 was determined based on a power analysis using effect sizes extracted from three closely related masked priming ERP studies (Carreiras et al., 2009; Holcomb et al., 2024; Y. Lee et al., 2016). Individual estimates ranged from *d* = 0.35 to *d* = 0.90. The inverse-variance weighted average across five estimates derived from these studies was *d* = 0.545 (95% CI [0.378, 0.712]). A paired-*t* power analysis using this estimate indicated that *N* = 30 affords 82.2% power (α = .05, two-tailed), exceeding the conventional 80% threshold. Additionally, since our primary analysis employs a Bayesian linear mixed-effects model (LMM) with crossed random effects for participants and items and 40 trial-level observations per condition, effective sensitivity is further increased relative to participant-averaged analyses, consistent with recommendations for trial-level modeling (Baayen et al., 2008).

Data from one participant was excluded due to poor behavioral performance (< 50% accuracy), resulting in twenty-nine participants (16 female; age: 23.31 ± 2.65 years). All participants were financially compensated for their participation and gave written informed consent before participation, in accordance with a protocol approved by Institutional Review Board of Korea University (KUIRB-2024-0321-01).

### 2.2. Procedure

A visual masked priming lexical decision task was used (see **Figure 1**). First, a fixation point (“+”) appeared at the center of the screen for 500 ms, followed by a forward mask (“####”) presented at the same location for 500 ms. After the mask, a prime stimulus was briefly presented for 50 ms, and the target stimulus appeared immediately after prime offset. During target presentation, participants were instructed to decide whether the visual stimulus was a real word or not by pressing a designated response key. Participants were instructed to minimize eye blinks during stimulus presentation and response execution. After the target presentation, an inter-trial interval (ITI) was presented for 1000, 1100, or 1200 ms, during which a visual cue (> <) was displayed, instructing participants to blink if needed. Each participant completed 24 practice trials and proceeded to the main experiment only after achieving an accuracy of at least 70%. All visual stimuli were presented in white Malgun Gothic font (size 40) on a black background and were displayed on a 24-inch LG monitor, with responses collected using a Serial Response Box, controlled using E-Prime 3.0 Professional (Psychology Software Tools, Pittsburgh, PA, USA).

**Figure 1.**
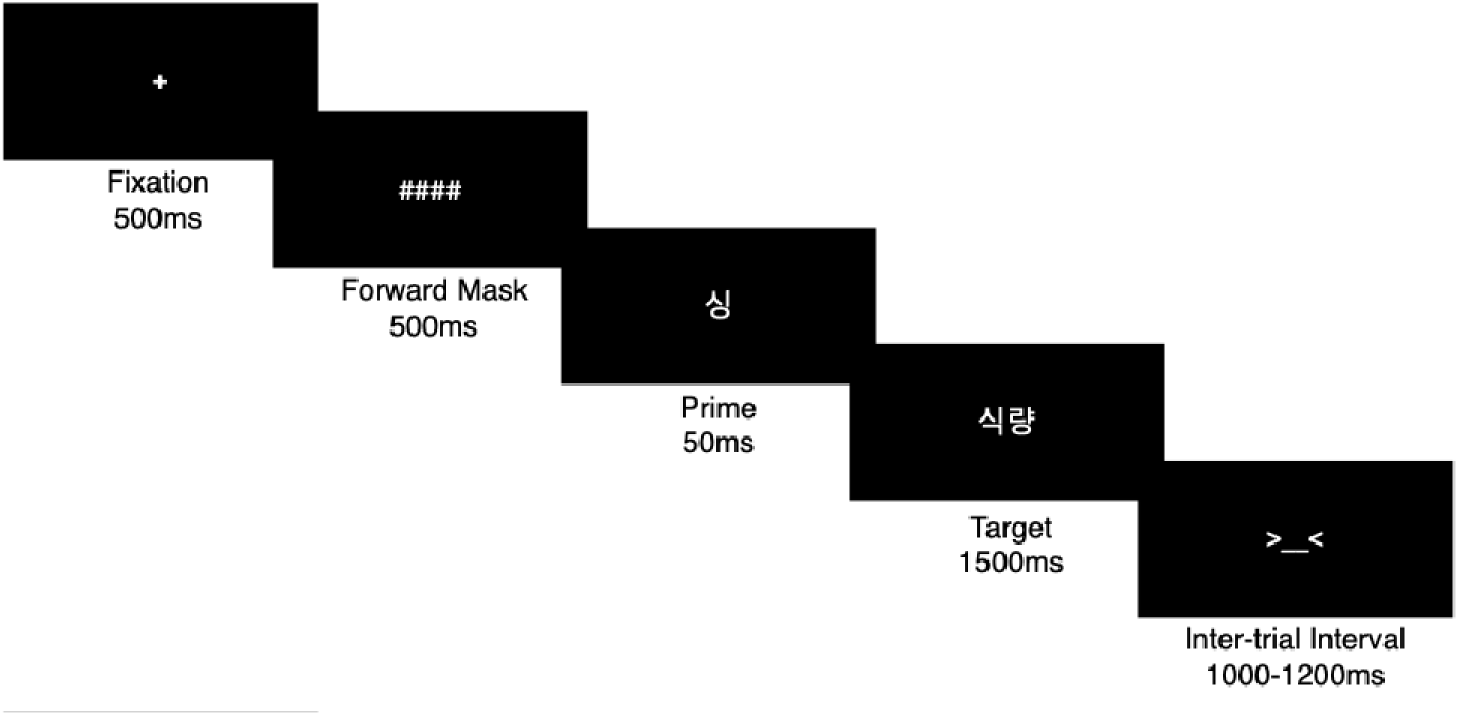
Schematic representation of the visual masked priming lexical decision task procedure.

### 2.3. Materials

A total of 120 Korean disyllabic words were selected as targets from the Korean Sejong corpus (15-million-word size; Kang & Kim, 2009). Every stimulus underwent phonological change at the first syllable to dissociate orthographic and phonological representations. Although Korean is known for its relatively consistent grapheme-to-phoneme mappings (Pae et al., 2020), phonological changes at the first syllable are not uncommon in Korean; a large-scale corpus analysis using the Sejong Corpus reported first-syllable phonological change rates of 9.21% for simple noun words and up to 20.93% for inflected verb forms (Kim et al., 2023). The mean Zipf frequency (i.e., log-transformed word occurrences per billion; van Heuven et al., 2014) of the word stimuli was 3.85 (*SD =* 0.50) and the mean log-transformed frequency of the first syllable of the target was 3.61 (*SD* = 0.52).

A Latin square design was used across stimulus lists to ensure that each target appeared once in each priming condition, to which ten participants were randomly assigned. Priming conditions were manipulated such that each list contained orthographically related (e.g., 식/ ik /-식량/ iŋ aŋ/), phonologically related (e.g., 싱/ iŋ/-식량/ iŋ aŋ/) and unrelated (e.g., 왼<u>/w n</u>/- 식량/<u>iŋ</u> aŋ/) conditions. Target stimuli were assigned to the three priming conditions, with 40 items per condition within a single list. Accordingly, each stimulus was presented once in the orthographically related, phonologically related, and unrelated conditions across the three lists. Finally, 120 disyllabic pseudowords were created by combining the first syllable of target words with another randomly selected syllable, serving as foils for the lexical decision task. The complete stimulus lists, along with the data and analysis code, are available on the Open Science Framework (https://osf.io/urybq/overview?view_only=02d7cb8a2eb141a98c3f9abdbba3593e).

### 2.4. EEG acquisition and preprocessing

Continuous EEG data were recorded using a 64-channel BrainAmp system (Brain Products GmbH, Munich, Germany) with 64 Ag/AgCl active electrodes. Electrodes were positioned according to the international 10-20 system using an actiCAP electrode cap (see **Figure 2A** for a detailed montage). Recordings were conducted in an electrically shielded, sound-attenuated room. All electrode impedances were maintained below 10 kΩ throughout the recording session. The online reference was set to FCz, and the ground electrode was positioned at FPz.

**Figure 2.**
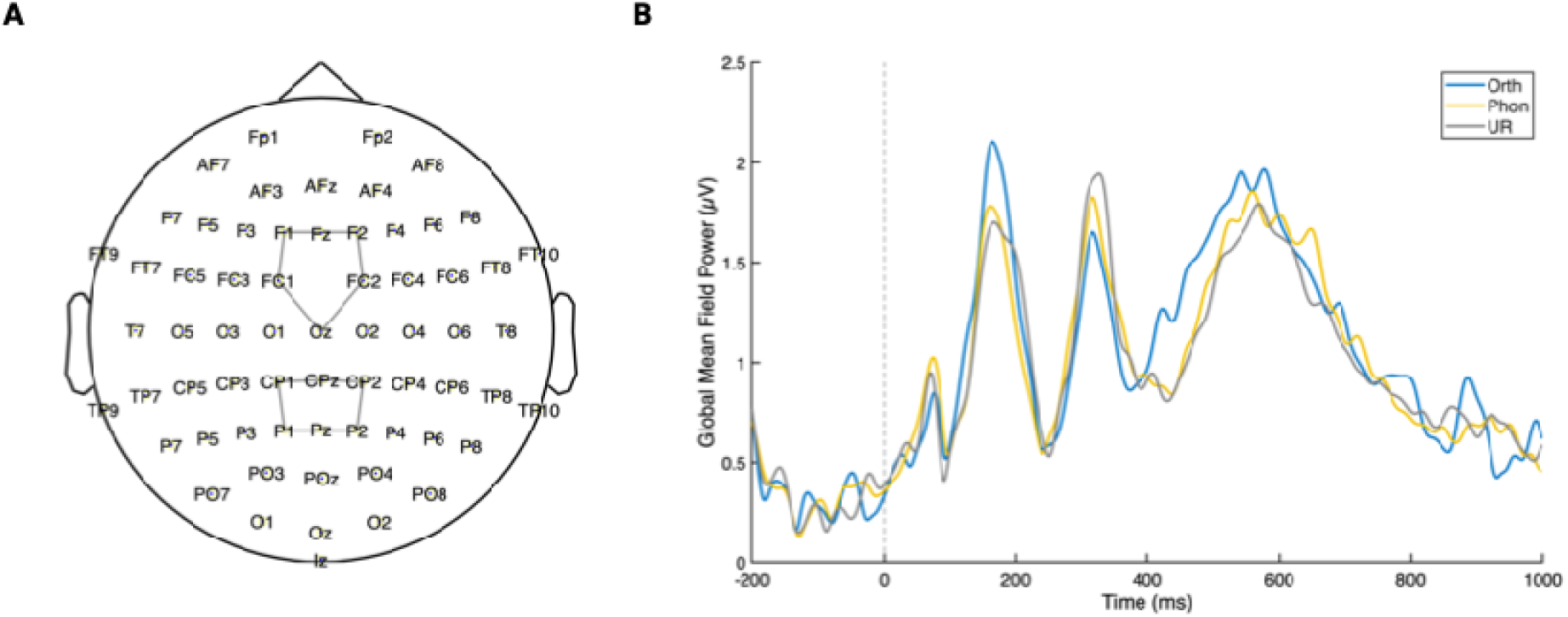
Electrode montage showing the 64-channel actiCAP electrode layout used for EEG recording according to the international 10-20 system. (B) Global mean field power across time for orthographic (Orth), phonological (Phon), and unrelated (UR) priming conditions, showing divergence in neural activity patterns across conditions from stimulus onset (0 ms) to 1000 ms post-stimulus.

All preprocessing procedures were conducted using a custom MATLAB script with EEGLAB (version 2025.1.0; Delorme & Makeig, 2004) implemented in MATLAB (The MathWorks, Inc., Natick, MA). Raw continuous EEG data were first downsampled to 500 Hz to reduce computational demands while preserving adequate temporal resolution for ERP analysis. A high-pass filter with a cutoff frequency of 0.1 Hz was applied using a FIR filter to remove slow drifts and DC offset while preserving low-frequency neural activity. Noisy channels were identified and removed using the clean_rawdata plugin from EEGLAB. Channels were flagged as noisy based on the following criteria: (1) flatline duration exceeding 5 seconds, (2) correlation with other channels below 0.8, and (3) line noise criterion of 4. The distance metric was set to Euclidean. Burst criterion, window criterion, and burst rejection were disabled during this step to focus exclusively on channel-level artifacts. The mean number of removed and interpolated channels was 3.24 (5.31% of all channels). Following bad channel removal, a low-pass filter with a cutoff frequency of 100 Hz was applied to attenuate high-frequency noise and electrical artifacts. Removed channels were then interpolated using spherical spline interpolation to restore the full 64-channel montage, ensuring consistent spatial sampling across all participants for subsequent analyses. Data were re-referenced to the common average reference by computing the average of all 64 channels at each time point and subtracting it from each channel.

Independent component analysis (ICA) was performed using the runica algorithm (Makeig et al., 1996) implemented in EEGLAB to decompose the EEG signal into maximally independent components. Independent components were then classified using the ICLabel plugin (Pion-Tonachini et al., 2019) with default settings. Components were marked for rejection based on the following criteria: (1) brain probability < 0.5, and (2) either muscle probability ≥ 0.8 or eye probability ≥ 0.6. The mean number of removed independent components was 5.31 (8.94% of all ICs). Identified artifact components were removed from the data by back-projecting the remaining components to channel space.

For ERP analysis, preprocessed continuous data underwent additional filtering with a bandpass of 1–30 Hz using pop_eegfiltnew function with default FIR filter settings. Data were then segmented into epochs ranging from –200 to 1000 ms relative to stimulus onset, with four experimental trigger codes marking the onset of experimental conditions (i.e., orthographically related, phonologically related, unrelated, and pseudowords). Baseline correction was performed by subtracting the mean amplitude of the –200 to 0 ms pre-stimulus interval from each epoch. Epochs were rejected if any channel exceeded an amplitude threshold of ±100 μV during the entire epoch window. Additionally, epochs corresponding to trials with incorrect behavioral responses or missing reaction times were excluded from analysis. The mean number of removed epochs was 29.93 (12.47% of all trials), and did not differ across experimental conditions [Orthographic: 5.86; Phonological: 5.52; Unrelated: 5.52; *F*(2, 56) = 0.248, *p* = .781]. Final ERP data consisted of correct-response, artifact-free epochs for each experimental condition.

For time-frequency analysis, a separate preprocessing stream was applied to the preprocessed continuous data. Data were filtered with a bandpass of 1–60 Hz to preserve oscillatory activity in relevant frequency bands. Epochs were extracted from –500 to 1500 ms relative to stimulus onset to provide adequate temporal padding for time-frequency decomposition. The same artifact and behavioral rejection criteria applied to ERP data were used to ensure consistency across analysis streams.

### 2.5. Behavioral analysis

Behavioral and ERP data were analyzed using Bayesian linear mixed-effects models (LMMs) using the brms package (Bürkner, 2017) in R (R Core Team, 2025), estimated via Hamiltonian Monte Carlo sampling (4 chains × 2,000 post-warmup iterations). All models included crossed random intercepts for participants and items, and Condition was reference-coded with Unrelated as the baseline level.

Reaction times (correct trials only) were modeled using a Gaussian family on log - transformed RT; accuracy was modeled using a Bernoulli family with logit link. Both models included Condition (reference-coded, Unrelated as baseline) and List as fixed factors with crossed random intercepts for participants and items. Weakly informative priors were placed on fixed effects: Normal(0, 0.1) on the log scale for RT and Normal(0, 1) on the logit scale for accuracy.

We report the evidence ratio for one-sided hypotheses (Evid.Ratio = pd/(1–pd), where pd is the posterior probability of direction that the effect is in the predicted direction), which quantifies how much more probable the effect is in one direction than the other. When the 95% credible interval (CI) included zero, we additionally report the evidence ratio for the point null hypothesis (Evid.Ratio ) computed via the Savage-Dickey density ratio method, equivalent to a Bayes factor (Wagenmakers et al., 2010). For Evid.Ratio , we follow the interpretation scale proposed by M. D. Lee and Wagenmakers (2012): values of 1–3, 3–10, 10–30, and >30 indicate anecdotal, moderate, strong, and very strong evidence for the null, respectively. Evid.Ratio values are interpreted continuously as the odds favoring one direction over the other; values ≥ 19 (corresponding to pd ≥ .95) indicate strong directional evidence.

### 2.6. ERP analysis

ERP time windows were pre-defined based on previous literature on syllable processing (Barber et al., 2004; Hutzler et al., 2004; Y. Kwon & Lee, 2015) and visual inspection of the global field power (see **Figure 2B**): P200 (150–250 ms) and N400 (350–550 ms). For each ERP component, amplitudes were extracted and averaged over two regions of interest (ROIs; **Figure 2A**): fronto-central (Fz, F1, F2, FC1, FC2, Cz) and centro-parietal sites (CPz, CP1, CP2, P1, P2, Pz).

Separate Bayesian LMMs were fitted to single-trial ERP amplitudes for each combination of component (P200: 150–250 ms; N400: 350–550 ms) and ROI (fronto-central, centro-parietal), yielding four primary models. Each model included Condition (reference-coded, Unrelated as baseline) and List as fixed factors, with crossed random intercepts for participants and items. A weakly informative prior of Normal(0, 1) on the μV scale was placed on all fixed effects. Convergence was confirmed by R = 1.00 for all parameters. In addition, full models combining both ROIs — with ROI (sum-coded), Condition, their interaction, and List as fixed factors — were fitted for each component as confirmatory analyses; these results are reported in Supplementary Materials (**Tables S1** and **S4**).

To complement the hypothesis-driven ROI analysis, exploratory cluster-based permutation tests (CBPT) were conducted using FieldTrip toolbox (Oostenveld et al., 2011) implemented in MATLAB. Spatial neighborhood structure was defined using triangulation method on the actiCAP64 electrode layout. The analysis tested for differences between each pair of experimental conditions (Orthographic vs. Unrelated, Phonological vs. Unrelated,

Orthographic vs. Phonological) across all time points and channels. The cluster-forming threshold was set at α = 0.05 (two-tailed), with a minimum of 2 neighboring channels required to form a cluster. Cluster-level statistics were computed using the maxsum method, and significance was determined through Monte Carlo permutation (10000 randomizations) with a corrected α = 0.025 (two-tailed) to control for Type I error rate. The dependent-samples t-statistic was used as the test statistic, with subjects as the unit of observation.

### 2.7. TFR analysis

Time-frequency analysis was conducted using the FieldTrip toolbox (Oostenveld et al., 2011) implemented in MATLAB to examine oscillatory dynamics in response to experimental conditions. Preprocessed EEG data (filtered 1–60 Hz, epoched –500 to 1500 ms) were converted from EEGLAB to FieldTrip format for analysis. Time-frequency representations (TFRs) were computed using Morlet wavelet convolution. Frequencies of interest ranged from 2 to 30 Hz in 1 Hz steps. The number of wavelet cycles increased linearly with frequency (cycles = frequency / 2), providing an optimal trade-off between temporal and frequency resolution across the frequency range. Power estimates were computed at 10 ms intervals (time of interest: –500 to 1500 ms). Baseline correction was applied using the relative change method [(power – baseline) / baseline] with a baseline window of –200 to 0 ms, consistent with ERP analysis.

To examine frequency-specific effects, TFR data were analyzed within pre-defined frequency bands of interest: theta (4–8 Hz), lower beta (13–20 Hz), and upper beta (20–30 Hz). Single-trial power estimates were averaged within each frequency band for subsequent statistical analysis.

Cluster-based permutation tests were conducted separately for each frequency band to identify spatiotemporal clusters showing significant condition differences while controlling for multiple comparisons across time points and channels. The analysis compared each pair of experimental conditions (Orthographic vs. Unrelated, Phonological vs. Unrelated, Orthographic vs. Phonological) using the same CBPT parameters as described for ERP analysis. Spatial neighborhood structure was defined using the triangulation method on the 64-channel layout.

Statistical testing was performed within the time window of –200 to 1000 ms. The cluster-forming threshold was set at α = 0.05 (two-tailed), with a minimum of 2 neighboring channels required to form a cluster. Cluster-level statistics were computed using the maxsum method, and significance was determined through Monte Carlo permutation (10000 randomizations) with a corrected α = 0.025 (two-tailed). The dependent-samples *t*-statistic was used, with subjects as the unit of observation. This analysis tested for condition differences within specific frequency ranges while accounting for the spatiotemporal extent of oscillatory effects.

## 3. RESULTS

### 3.1. Behavioral results

Overall, participants maintained high accuracy across all priming conditions (> 90.6%) with mean reaction times of 612 ms. **Figure 3** represents the means and standard errors of the RTs and accuracy for two related (i.e., orthographic and phonological) and unrelated priming conditions.

**Figure 3.**
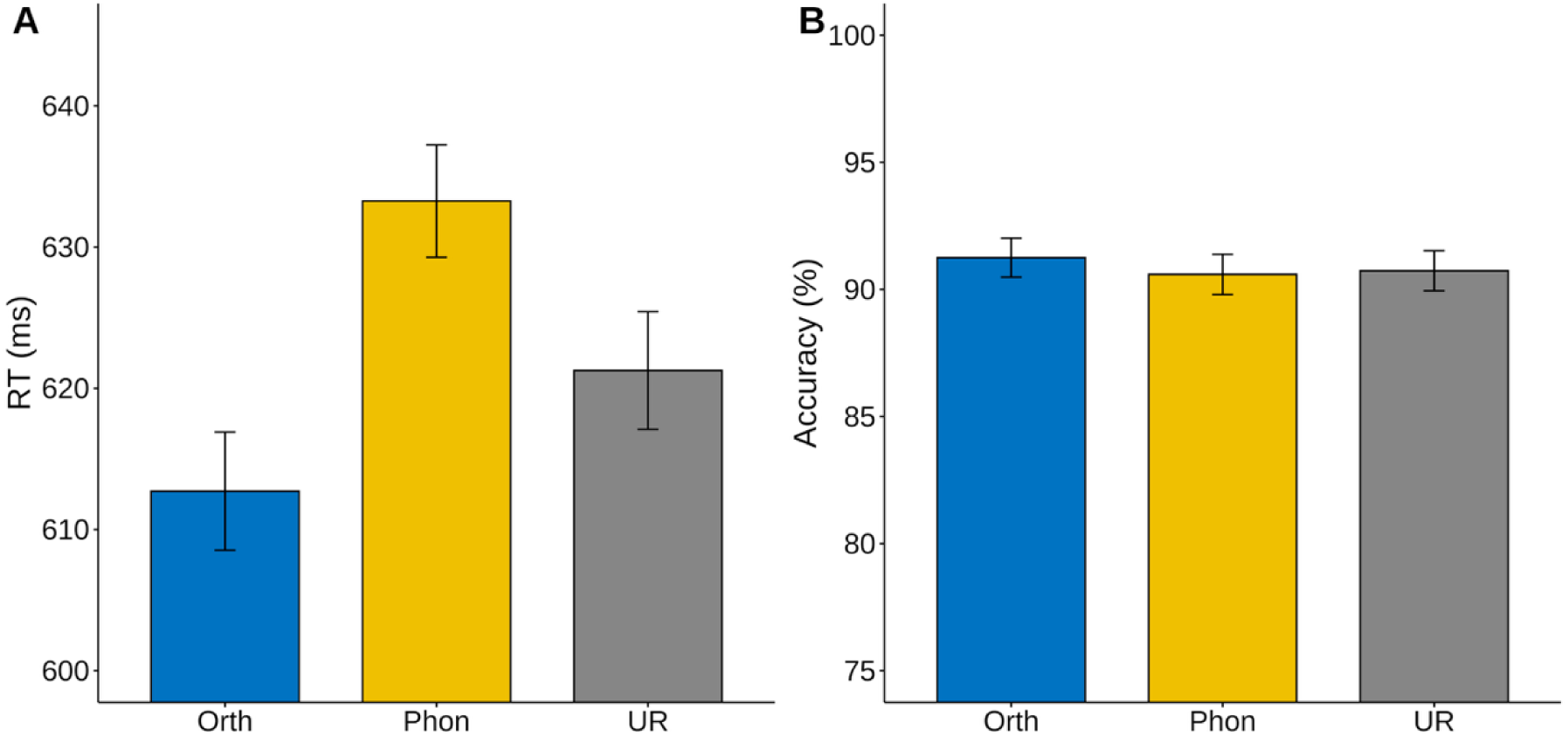
Behavioral performance in the masked priming lexical decision task (A) Mean reaction times (ms) for orthographic, phonological, and unrelated priming conditions. Error bars represent standard error. Orthographic priming produced significantly faster responses than both phonological and unrelated conditions. (B) Mean accuracy rates (%) across the three priming conditions. High accuracy was maintained across all conditions (>90%) with no significant differences between conditions.

Bayesian LMMs of log-transformed reaction times revealed strong evidence for an orthographic facilitation effect (Orthographic vs. Unrelated: β = –0.011, 95% CI [–0.017, – 0.006], Evid.Ratio = 7,999) and a phonological inhibition effect (Phonological vs. Unrelated: β = 0.006, 95% CI [0.001, 0.012], Evid.Ratio = 71.1). Orthographic primes produced substantially faster responses than phonological primes (Evid.Ratio > 39,999). The List effect showed no evidence of an effect across both models (Evid.Ratio < 3).

The Bayesian LMM for accuracy revealed no reliable condition differences.

Orthographic primes did not differ from unrelated primes (β = 0.191, 95% CI [−0.144, 0.525], Evid.Ratio = 3.35), nor did phonological primes differ from unrelated primes (β = −0.083, 95% CI [−0.418, 0.248], Evid.Ratio = 5.45). The orthographic versus phonological comparison yielded only anecdotal evidence for the null (β = 0.274, 95% CI [−0.064, 0.610], Evid.Ratio = 2.43).

The behavioral results demonstrated a clear orthographic facilitation effect in reaction times, with faster responses following orthographically related primes compared to both unrelated and phonologically related primes. In contrast, phonological priming produced reliably slower responses than the unrelated baseline (Evid.Ratio = 71.1), indicating a phonological inhibition effect rather than facilitation. Accuracy remained high across all conditions (> 90%), with no meaningful condition differences (all Evid.Ratio = 2.33–5.37), confirming that RT effects reflect priming rather than speed-accuracy trade-offs.

### 3.2. ERP results

Grand-average ERPs for the three priming conditions are presented in **Figure 4**, showing waveforms at fronto-central (**Figure 4A**) and centro-parietal (**Figure 4B**) regions of interest, along with topographic distributions of the difference waves (**Figure 4C**). ERP amplitudes were analyzed using Bayesian LMMs — separately for each component (P200: 150–250 ms; N400: 350–550 ms) and ROI (FrontoCentral, CentroParietal). Full models combining both ROIs are provided in **Supplementary Results (Tables S1** and **S4**); all ROI-specific model outputs are provided in **Tables S2–S3** (P200) and **S5–S6** (N400).

**Figure 4.**
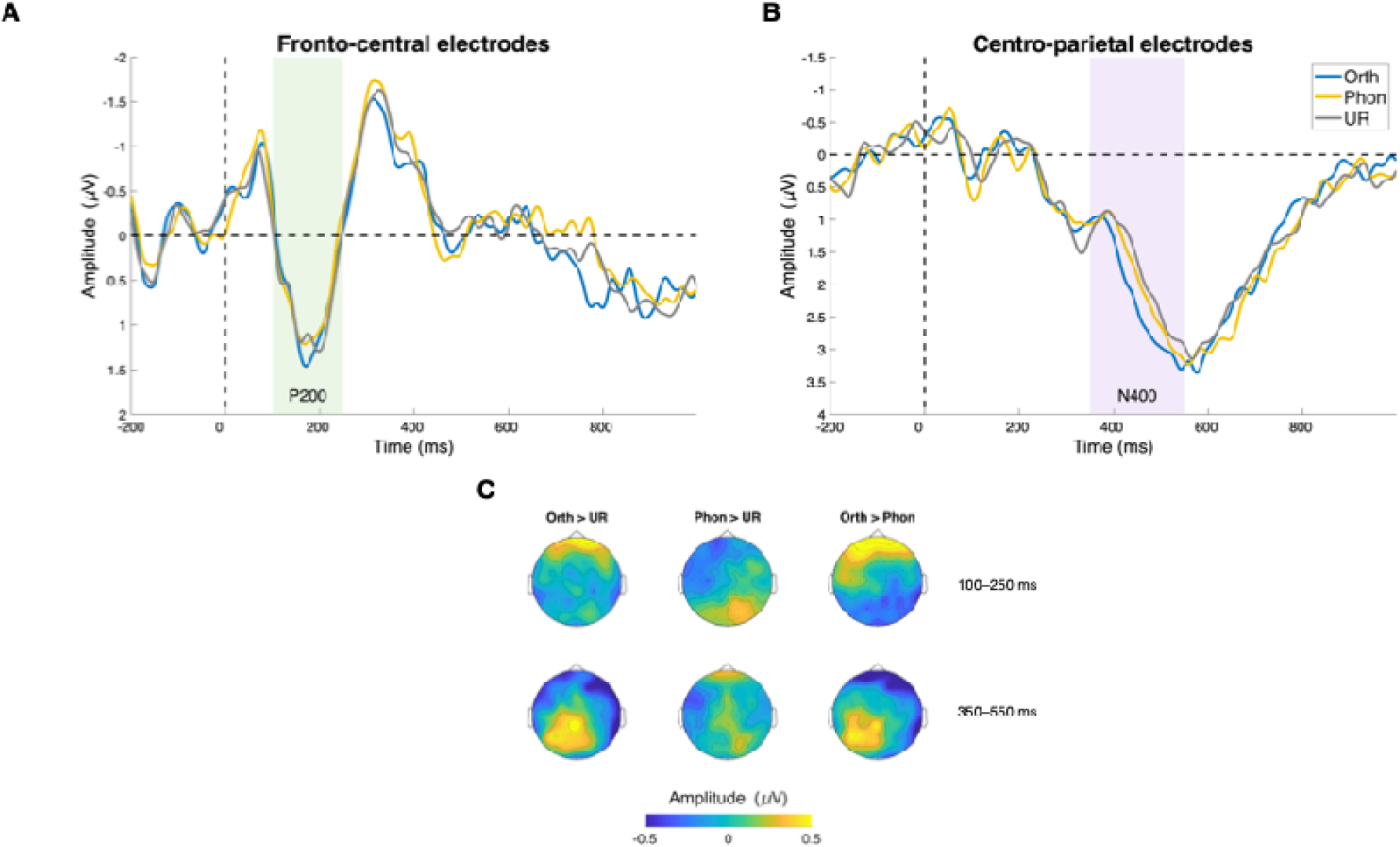
(A) Grand-averaged ERP waveforms at fronto-central electrodes (Fz, F1, F2, FC1, FC2, Cz) showing the P200 component (150–250 ms, highlighted in green). (B) Grand-averaged ERP waveforms at centro-parietal electrodes (CPz, CP1, CP2, P1, P2, Pz) showing the N400 component (350–550 ms, highlighted in purple). Negative amplitudes are plotted upwards. (C) Topographic distributions of difference waves for orthographic versus unrelated, phonological versus unrelated, and orthographic versus phonological comparisons in the P200 (150–250 ms) and N400 (350–550 ms) time windows. Color scale represents amplitude in µV.

#### 3.2.1. P200 (150–250 ms)

Bayesian LMMs at fronto-central sites (**Figures 4A and 4C, Table S2**) revealed larger P200 amplitudes for orthographic than unrelated primes (β = 0.342 μV, 95% CI [0.069, 0.616], Evid.Ratio = 140.34). No reliable differences were observed for phonological versus unrelated primes at fronto-central sites (β = 0.151 μV, 95% CI [−0.125, 0.424], Evid.Ratio = 3.96; **Table S2**). At centro-parietal sites, no condition showed evidence of P200 modulation (all Evid.Ratio < 5, all 95% CIs including zero; **Table S3**).

#### 3.2.2. N400 (350–550 ms)

Bayesian LMMs revealed a clear centro-parietal pattern (**Figures 4B–C**, **Table S5**). At centro-parietal sites, orthographic primes produced a robust N400 amplitude reduction (more positive; β = 0.317 μV, 95% CI [0.059, 0.576], Evid.Ratio = 116.6), and differed substantially from phonological primes (β = 0.298 μV, 95% CI [0.040, 0.558], Evid.Ratio = 81.0). In contrast, phonological primes did not differ from unrelated primes at centro-parietal sites (β = 0.019 μV, 95% CI [−0.238, 0.277], Evid.Ratio = 7.73). At fronto-central sites, no reliable N400 modulation was observed for either orthographic or phonological priming (all 95% CIs including zero; Evid.Ratio = 6.30–7.44; **Table S6**).

#### 3.2.3. Cluster-Based Permutation Tests

To complement the hypothesis-driven ROI analysis, exploratory cluster-based permutation tests (CBPT) were conducted to identify spatiotemporal clusters showing significant condition differences across all time points and channels. CBPT results are displayed in **Figure 5**, showing spatiotemporal distributions of significant effects across all channels and time points.

**Figure 5.**
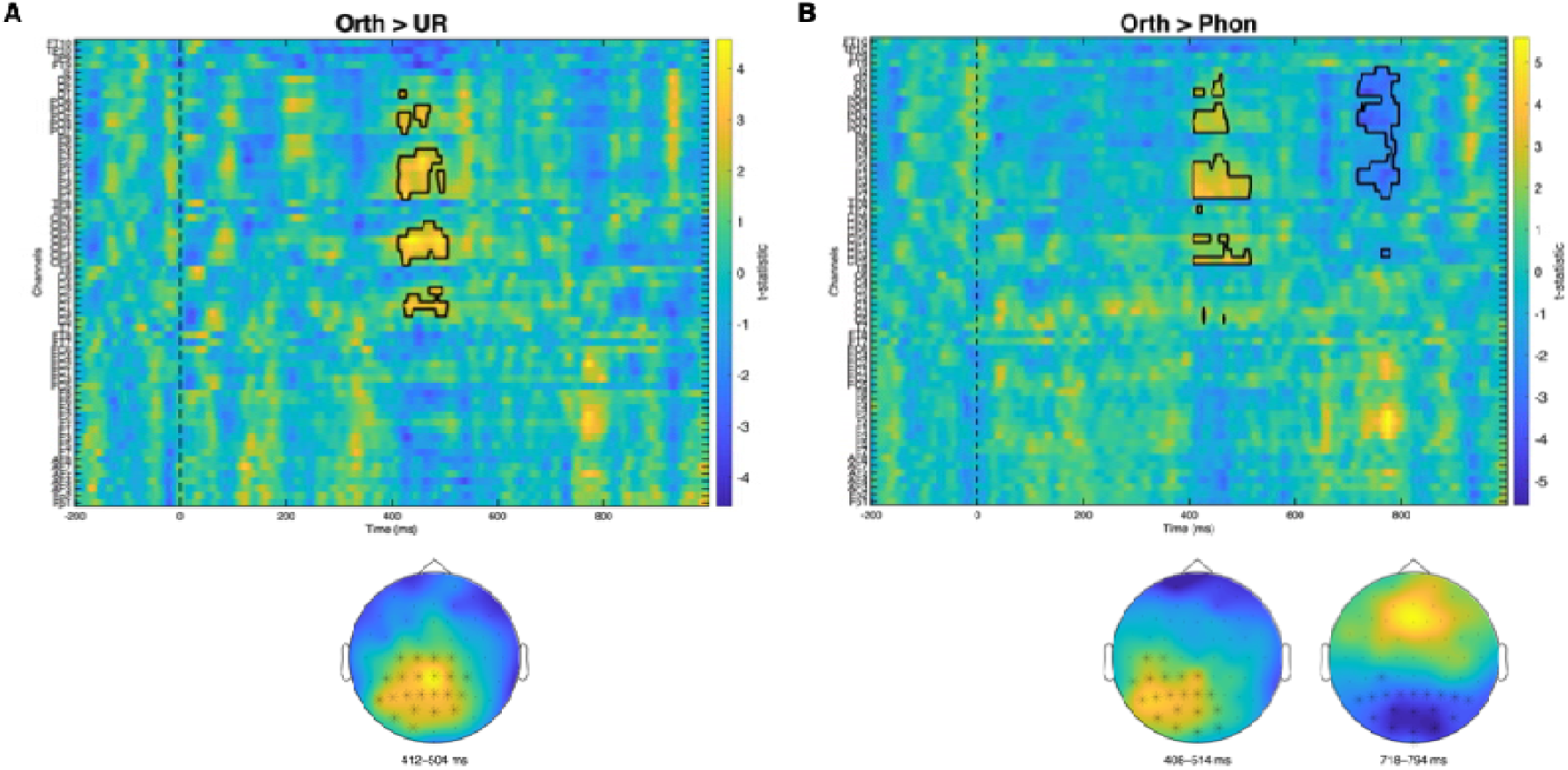
Cluster-based permutation test results for ERP contrasts (A) Orthographic versus unrelated contrast showing a positive cluster (412–504 ms, cluster statistics (sum of *t*-values within cluster) = 1495.9, *p* = .0142) across centro-parietal regions. Channel-by-time representation with significant clusters outlined in black (top) and topographic distribution of the cluster at 412–504 ms (bottom). (B) Orthographic versus phonological contrast showing a positive cluster (408–794 ms, cluster statistic = 1257.9, *p* = .0119) and a negative cluster (718–794 ms, cluster statistic = –1120.6, *p* = .0171) across centro-parietal and posterior regions. Channel-by-time representation with significant clusters outlined in black (top) and topographic distributions at early (408–514 ms) and late (718–794 ms) phases of the cluster (bottom). Color scale represents t-statistic values.

CBPT revealed a significant positive cluster for orthographic versus unrelated conditions (**Figure 5A**; 412–504 ms, *p* = .0142) distributed across 21 centro-parietal and posterior electrodes (C2, C3, CP1, CP2, CP3, CP4, CP5, CPz, Cz, O1, P1, P2, P3, P4, P5, P7, PO3, PO4, PO7, POz, Pz). This cluster overlaps temporally with the N400 window and reflects sustained orthographic facilitation with a broad centro-posterior distribution.

There was also a significant positive cluster for orthographic versus phonological conditions (**Figure 5B**; 408–514 ms, *p* = .0119) spanning 20 centro-parietal and posterior electrodes (C3, C5, CP1, CP3, CP5, CPz, O1, O2, Oz, P1, P2, P3, P5, P7, PO3, PO4, PO7, POz, Pz, TP7). This extended cluster suggests prolonged differential processing between orthographic and phonological priming, extending well beyond the N400 window into late posterior positivity. Notably, within the later portion of this interval, a significant negative cluster emerged (718–794 ms, *p* = .0171), encompassing 19 posterior parieto-occipital electrodes (P1, P2, P3, P4, P5, P6, P7, P8, Pz, PO3, PO4, PO7, PO8, POz, O1, O2, Oz, Iz, and CP3). The negative polarity of this cluster reflects reduced late positivity for orthographic relative to phonological priming, following the earlier reduced N400 amplitude observed for orthographic priming in the same contrast. Together, these effects suggest sustained orthographic facilitation from the N400 window into later processing stages. No significant clusters emerged for phonological versus unrelated conditions, consistent with the absence of reliable phonological ERP modulation in the ROI-based analyses.

### 3.3. Time-frequency results

Time-frequency analysis examined oscillatory dynamics within predefined frequency bands (theta: 4–8 Hz, lower beta: 13–20 Hz, upper beta: 20–30 Hz) using cluster-based permutation testing (two-tailed, *p* < 0.025, 10000 randomizations). Significant clusters are displayed in **Figure 6**.

**Figure 6.**
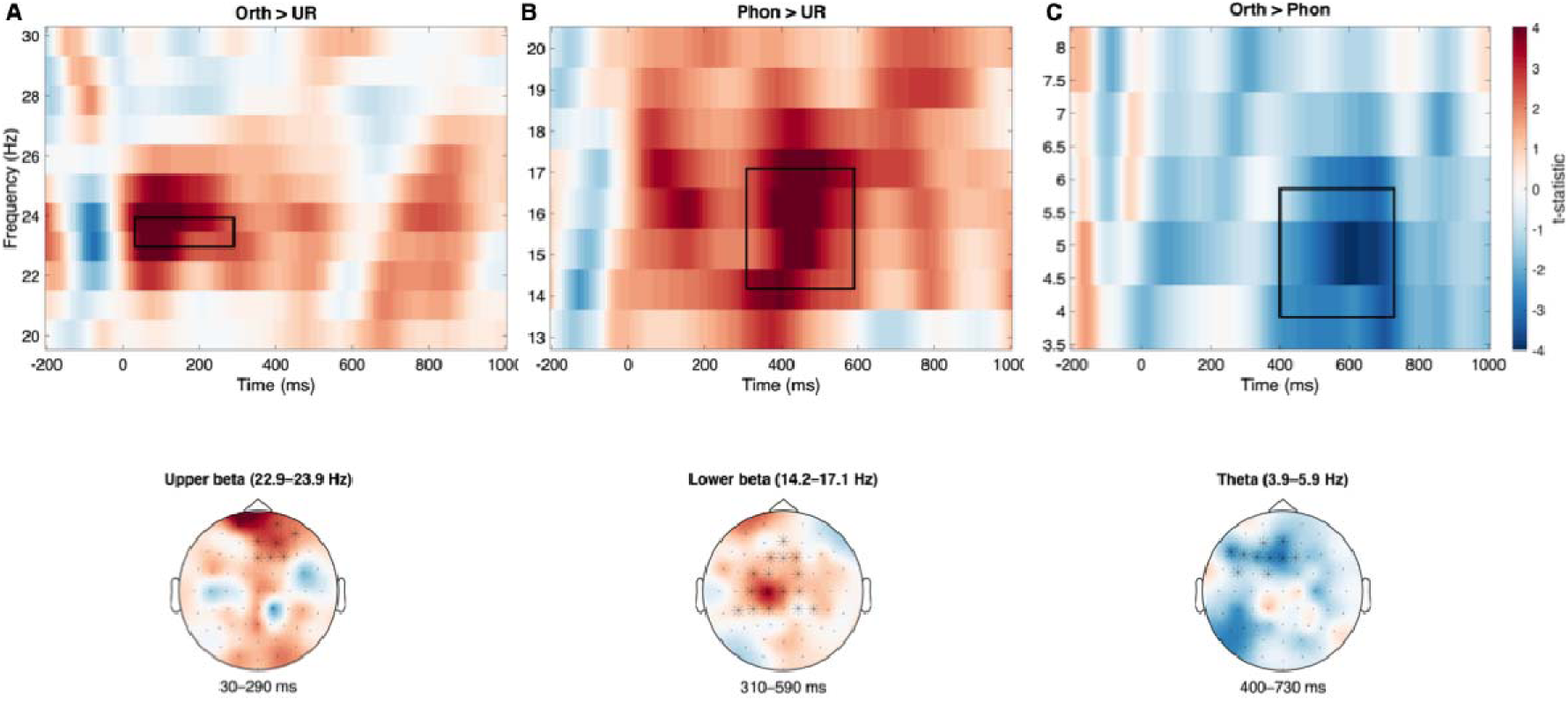
Time-frequency representations (TFRs) showing significant clusters identified using cluster-based permutation testing. (A) Orthographic versus unrelated contrast (Orth > UR) showing early upper-beta synchronization (22.9–23.9 Hz, 30–290 ms; *p* = .0243; cluster statistic (sum of t-values within cluster) = 574.79) over frontal electrodes, displayed at the peak channel Fp1. (B) Phonological versus unrelated contrast (Phon > UR) showing sustained lower-beta synchronization (14.2–17.1 Hz, 310–590 ms; *p* = .0128; cluster statistic (Σt) = 855.36) over fronto-central electrodes, displayed at the peak channel C1. (C) Orthographic versus phonological contrast (Orth > Phon) showing reduced theta power (3.9–5.9 Hz, 400–730 ms; *p* = .0211; cluster statistic (Σt) = −825.74) for orthographic priming over fronto-central electrodes, displayed at the peak channel Fz.

#### 3.3.1. Upper Beta Band (20–30 Hz)

Orthographically related words elicited a significantly enhanced upper beta band power relative to the unrelated condition in a positive cluster (*p* = .0243; **Figure 6A**) spanning 30–290 ms (22.9–23.9 Hz) across eight frontal electrodes (Fp1, Fz, F4, Fp2, AFz, AF8, AF4, F2). The peak effect occurred at Fp1: *t* = 4.57, 90 ms). This early frontal beta synchronization temporally overlapped with the P200 window and parallels the orthographic P200 enhancement observed in the ERP analysis, which may suggest an early orthographic encoding process.

#### 3.3.2. Lower Beta Band (13–20 Hz)

Phonological priming elicited a significant positive cluster (*p* = .0128; **Figure 6B**) spanning 310–590 ms (14.2–17.1 Hz) across sixteen fronto-central and central electrodes (Fz, FC1, C3, CP5, CP1, CP2, Cz, FC2, AFz, F1, FC3, C1, CP3, CP4, C2, F2). The peak effect was observed at C1 (450 ms, 16.1 Hz; *t* = 4.54). This sustained central beta increase in the mid-latency window suggests enhanced processing demands for phonological priming during lexical access. The temporal profile of this effect extends beyond the P200 window but overlaps with the early portion of the N400 time window, indicating that phonological priming engages prolonged beta-band oscillatory activity that may reflect ongoing lexical competition or phonological assembly processes.

#### 3.3.3. Theta Band (4-8 Hz)

In a later time window, orthographic priming was associated with reduced theta power relative to phonological priming (*p* = .0211; **Figure 6C**) between 400 and 730 ms (3.9–5.9 Hz) across ten fronto-central electrodes (Fz, F3, F7, FC5, FC1, AF3, AFz, F1, F5, F2). The peak effect occurred at Fz (*t* = –3.95, 600 ms). This theta reduction overlapped temporally with the N400 window, during which orthographic priming produced robust centro-parietal N400 reductions. Reduced frontal theta power for orthographic priming overlapped with the N400 window, consistent with reduced processing demands during lexical access.

In sum, the TFR results demonstrated early upper beta synchronization for orthographic priming, sustained lower beta increase for phonological priming, and later theta suppression for orthographic versus phonological priming.

## 4. Discussion

The present study employed masked priming with lexical decision and high-density EEG to simultaneously address two questions regarding sublexical processing in Korean visual word recognition: (1) whether syllabic priming effects reflect orthographic or phonological syllable overlap, and (2) whether these effects originate at early form-encoding stages or at later lexical stages during competitive activation. Exploiting Korean alpha-syllabic structure, in which morphophonemic spelling principles allow systematic dissociation of orthographic from phonological syllable representations, we contrasted orthographically identical and phonologically overlapping primes while recording both event-related potentials and time-frequency representations. Our findings revealed a clear spatiotemporal dissociation: orthographic priming produced early fronto-central P200 enhancement (150–250 ms) accompanied by upper beta synchronization (30–290 ms) and subsequent centro-parietal N400 reduction (350–550 ms), along with behavioral facilitation. Phonological priming, by contrast, produced no early electrophysiological modulation and produced behavioral inhibition rather than facilitation, while eliciting sustained lower beta activity (310–590 ms) over central regions. These results are more consistent with a sequential or cascaded account of orthographic-phonological coordination at the syllable level, as predicted by dual-route models (Coltheart et al., 2001; Perry et al., 2007), than with fully parallel activation frameworks (Diependaele et al., 2010; Grainger & Ziegler, 2011).

### 4.1. Early Orthographic Processing: P200, N/P150, and Upper Beta Synchronization

Consistent with our prediction that orthographic syllable overlap would facilitate early visual-orthographic encoding, orthographically identical primes produced robust early electrophysiological modulation. The P200 component (150–250 ms) showed enhanced amplitude for orthographic primes over fronto-central sites, with no comparable modulation for phonological primes. It should be noted, however, that Korean lacks a case distinction, so the orthographic priming condition involved physical identity between the prime and the first syllable of the target. The early ERP effects observed here could therefore reflect, at least in part, low-level visual repetition rather than purely abstract orthographic processing. With this caveat in mind, the pattern nonetheless addresses the first research question by suggesting that the early ERP effect is driven by visual-orthographic overlap rather than phonological syllable representations. The crossed design, in which orthographic and phonological overlap were independently manipulated, resolves an ambiguity that has persisted in the alphabetic SFE literature, where P200 modulations by syllable frequency could not be attributed to either code in isolation (Barber et al., 2004; Hutzler et al., 2004).

The present P200 effect can be situated within the temporal benchmark of early ERP components in visual word recognition established by Grainger et al. (2006; see also Grainger & Holcomb, 2009). In their masked priming work, the N/P150 component (approximately 100–200 ms) was identified as an index of early sublexical form processing, exhibiting enhanced positivity at frontal sites and enhanced negativity at posterior sites when prime and target share visual-orthographic features—hence the N/P150 designation. It is important to note that this component taxonomy was developed within a parallel activation framework (the BIAM; Grainger & Ziegler, 2011), in which both orthographic and phonological codes are assumed to be activated rapidly and simultaneously. However, even within this framework, the N/P150 is characterized as primarily sensitive to visual-orthographic overlap rather than phonological overlap, reflecting the initial stages of letter and sub-word representation formation prior to phonological engagement (Carreiras et al., 2009; Grainger et al., 2006; Holcomb & Grainger, 2006).

The P200 enhancement observed in the present study aligns temporally and topographically with the early form-processing component described in parallel activation frameworks (Grainger & Holcomb, 2009). Crucially, while our experimental design adopted the same morphophonemic manipulation as Lee et al. (2016)—contrasting orthographic identity with phonological overlap—the resulting ERP patterns revealed a clearer functional dissociation. Lee et al. (2016) reported that both orthographic and phonological primes modulated the N/P150 component, suggesting a broad sensitivity to sublexical overlap. In contrast, the present study found a P200 enhancement that was selective for orthographic primes, with no evidence of significant phonological modulation in this time window. While Lee et al. (2016) observed no significant priming effects in their N250 window (300–400 ms), our study identified a robust orthographic facilitation effect in the N400 window (350–550 ms), coupled with significant behavioral priming. This implies that the previously reported N250 null effects might have stemmed from a window that only partially overlapped with lexical-level processing. Consequently, our results are consistent with a temporal pattern in which form-based orthographic analysis (P200) temporally precedes lexical-semantic integration (N400), with no evidence of early phonological activation in the present paradigm.

The absence of phonological priming effects at later latencies is particularly informative when considered against the N250 component. The N250 (approximately 200–350 ms) has been interpreted to index grapheme-phoneme conversion, reflecting the mapping from orthographic to phonological representations (e.g, Grainger et al., 2006). In alphabetic masked priming studies, phonologically overlapping primes that are orthographically dissimilar (e.g., pseudohomophone primes) typically modulate the N250, indicating that phonological codes become available at this processing stage. Under the parallel activation account (i.e., the BIAM; Grainger & Ziegler, 2011), phonological primes sharing sublexical phonological overlap should produce N250 modulation reflecting rapid and automatic phonological code activation. However, this prediction was not borne out in the present data: phonological primes produced neither early N/P150–P200 modulation nor later N250-range effects. This null result converges with Lee et al. (2016), who reported that while both orthographic and phonological primes modulated the N/P150 relative to unrelated primes, neither condition produced N250 effects (300–400 ms). The authors interpreted the results as a limited role of phonological information in early Korean visual word recognition. The absence of N250 modulation across both studies places constraints on the BIAM’s prediction that phonological codes are activated rapidly and in parallel with orthographic codes, instead supporting a sequential account in which orthographic processing achieves functional priority before phonological assembly begins. This temporal ordering is more consistent with the architecture of the DRC model, in which the direct orthographic route operates faster than the indirect phonological route (Coltheart et al., 2001).

The concurrent upper beta synchronization (22.9–23.9 Hz, 30–290 ms) over frontal regions provides insight into the computational mechanism underlying this rapid orthographic encoding. Upper beta synchronization has been broadly implicated in top-down processing and endogenous content (re)activation—the internally driven formation of cortical representations in the service of current task demands (Engel & Fries, 2010; Spitzer & Haegens, 2017). In the domain of language, beta oscillations are particularly suited for long-range cortical communication, supporting integrative functions across distant brain areas during word and sentence processing (Weiss & Mueller, 2012).

In the context of Hangul, where graphemes are geometrically arranged into compact syllable blocks, this early frontal upper beta burst likely reflects top-down driven integration of graphemic features into holistic syllable units, consistent with the role of beta synchronization in maintaining and reactivating stored orthographic representations (Lewis & Bastiaansen, 2015; Spitzer & Haegens, 2017). Notably, beta oscillations have been shown to correlate with the N400 component during language comprehension (Wang et al., 2012), and to index lexical frequency effects during word production (Piai et al., 2014), further suggesting lexical access mechanisms associated with the beta band. The convergence of P200 enhancement and upper beta synchronization in the same early time window provides complementary evidence, where the ERP effect indexes enhanced neural responsivity to familiar orthographic patterns. The oscillatory signature may reflect a top-down stabilization process that supports the consolidation of these patterns, though the precise functional role of upper beta synchronization in this context warrants further investigation. Together, these findings indicate that the alpha-syllabic structure of Korean enables efficient visuospatial parsing of syllable units during the initial sweep of visual processing, consistent with the early locus of orthographic effects reported in alphabetic languages (Ferrand & Grainger, 1992, 1993).

### 4.2. Phonological Processing: Lower Beta Dynamics and Behavioral Inhibition

The phonological condition revealed a pattern that addresses both research questions. Phonological primes produced no modulation of either P200 or N400 amplitudes, yet they elicited behavioral inhibition and sustained lower beta power increase (14.2–17.1 Hz, 310–590 ms) over central regions. This pattern indicates that phonological syllable overlap engages neural processes that are distinct from, and temporally delayed relative to, orthographic encoding—confirming the representational dissociation—while the absence of early modulation and the presence of behavioral inhibition constrains the processing stage at which these effects operate.

Lower beta activity (13–20 Hz) has been associated with maintenance of cognitive states and effortful processing in both domain-general (Engel & Fries, 2010) and language-specific contexts (Klimesch et al., 2001; Mai et al., 2016; Weiss & Mueller, 2012). Specifically, beta synchronization has been associated with binding mechanisms, with the lower beta band reflecting top-down integration of semantic, pragmatic, and syntactic information (Weiss & Mueller, 2012). In the context of visual word recognition, Klimesch et al. (2001) specifically identified that lower beta (beta-1b) dynamics over left language areas reflect graphemic-phonetic encoding, linking this frequency band to the translation of orthographic codes into phonological forms. Complementing this, Mai et al. (2016) reported that, during auditory sentence processing, lower beta (13–20 Hz) dynamics index the retrieval of stored phonological information and the maintenance of phonological units in working memory. More broadly, beta oscillations have been linked to the maintenance of current linguistic representations and top-down predictive processing (Lewis et al., 2015; Lewis & Bastiaansen, 2015).

One possible interpretation of the sustained lower beta increase is that it reflects the joint contribution of two processes. First, phonological primes activated shared phonological representations (/ iŋ/), requiring assembly of phonological codes from orthographically distinct visual input (‘싱’ vs. ‘식’). This assembly process is computationally more demanding than direct orthographic matching, as it requires grapheme-to-phoneme conversion and syllabic parsing along what dual-route models characterize as the indirect phonological route (Coltheart et al., 2001; Perry et al., 2010). Second, the orthographic mismatch between prime and target created representational conflict: the phonological overlap activated the target’s phonological representation while the orthographic non-identity simultaneously failed to pre-activate the target’s lexical representation, resulting in a net inhibitory effect. The observed behavioral inhibition is consistent with lexical competition driven by phonological overlap (Grainger & Jacobs, 1996; Lupker & Colombo, 1994), in which the prime activated a set of phonologically similar lexical candidates that competed with the target during selection, producing slower latencies relative to the unrelated baseline. This pattern aligns with the inhibitory syllable frequency effects reported in the Korean literature (Choi et al., 2015; Y. Kwon, 2012) and with the lexical competition account formalized in computational models (Conrad et al., 2010).

The temporal profile of this lower beta effect (310–590 ms) further constrains the processing stage of phonological syllable effects. The onset at 310 ms falls well after the P200 window in which orthographic effects emerged, converging with the trajectory observed in alphabetic masked priming studies that controlled for orthographic overlap (Carreiras et al., 2009), where reliable phonological modulation emerged only after 350 ms. These results suggest that phonological assembly operates subsequent to initial orthographic encoding rather than in parallel with it. Furthermore, the overlap of this oscillatory activity with the early portion of the N400 window is consistent with processing at or near the lexical level, and the concurrent behavioral inhibition suggests that phonological overlap may engage competitive processes among co-activated lexical candidates. This interpretation is supported by findings that inhibitory syllabic priming is contingent on lexical status (Dominguez et al., 1997), though the temporal overlap alone does not definitively localize these effects to a single processing level.

Critically, our findings challenge the interpretation that Korean readers do not automatically activate phonological information (Park, 1996; Bae & Yi, 2010; Tae et al., 2015). Rather, phonological representations are activated, as evidenced by the lower beta response, but this activation is computationally costly when orthographic information conflicts with phonological input. The convergence of behavioral inhibition with sustained oscillatory activity—in the absence of early ERP modulation—highlights the importance of combining behavioral and neural measures, and indicates that the pattern of phonological priming in Korean reflects active lexical competition rather than a simple absence of phonological processing.

The absence of early phonological effects, however, warrants separate consideration, as it may reflect structural and orthographical properties of Korean. As most Korean nouns are disyllabic, with each syllable built from multiple sub-syllabic letters, this imposes a greater representational load for phonological assembly than in languages with many high-frequency monosyllabic words. A very brief masked prime (e.g., 50–100 ms) may be sufficient to saturate orthographic facilitation driven by Hangul’s visually salient block structure, while falling short of the temporal window needed for complete phonological code assembly. Moreover, Hangul’s morphophonemic spelling system introduces an additional computational step, in which surface phonological forms often diverge from their underlying representations due to pervasive assimilation and neutralization. As a result, readers may need to compute both the orthographic block structure and the morphophonemic base before stabilizing a phonological code. Together, these properties may delay the emergence of detectable phonological facilitation relative to languages with shorter words and more transparent surface phonology (Lukatela & Turvey, 1994a, 1994b), rather than implying that the underlying mechanism differs across writing systems.

### 4.3. Lexical-Semantic Integration: N400 Reduction and Theta Suppression

The N400 window (350–550 ms) provided further evidence for the orthographic processing advantage. Orthographic priming produced robust N400 amplitude reduction at centro-parietal sites, confirmed by both ROI-based analysis and cluster-based permutation testing (412–504 ms, *p* = .0142). This N400 reduction was accompanied by reduced frontal theta power (3.9–5.9 Hz, 400–730 ms) for orthographic relative to phonological priming. Considering theta synchronization is canonically associated with lexical retrieval and semantic integration effort (Bastiaansen et al., 2008; Klimesch, 1999), the suppression of theta power observed in our data is consistent with reduced processing demands during lexical-semantic integration. Specifically, the valid orthographic prime likely pre-activates the target’s lexical entry, minimizing the need for the sustained memory search and retrieval operations typically indexed by theta synchronization.

The centro-parietal distribution of the N400 effect, combined with the absence of reliable N400 modulation at fronto-central sites, is consistent with the canonical topography of the N400 as an index of lexical-semantic integration rather than form-level processing (Kutas & Federmeier, 2011). The topographic dissociation between the early fronto-central P200 effect and the later centro-parietal N400 effect further supports the interpretation that orthographic priming facilitates processing across two distinct stages: early form encoding and subsequent lexical-semantic access. One plausible account is that the reduced N400 amplitude reflects a downstream benefit of early orthographic facilitation, where the rapid stabilization of the visual word form, indexed by P200 enhancement, facilitates lexical access, thus reducing neural resources required for subsequent semantic integration.

The absence of N400 or theta modulation for phonological primes indicates that shared phonology alone, without orthographic support, provides negligible benefit for semantic access in skilled Korean readers. This contrasts with findings in shallow orthographies where phonological priming can facilitate semantic processing (Ashby et al., 2009), highlighting script-specific adaptations. In Korean, the morphophonemic spelling system preserves morphological boundaries in orthography even when phonological changes occur at morpheme boundaries (e.g., 식 / ik / → 식량 / iŋ aŋ/). This orthographic stability may reduce functional reliance on phonological recoding, allowing skilled readers to develop a visual-orthographic route optimized for direct lexical access.

### 4.4. Sequential Processing and Theoretical Implications

Taken together, the behavioral, ERP, and oscillatory evidence suggest a coherent temporal pattern that bears on both research questions. Orthographic syllable overlap produced a cascade of effects beginning with early form encoding (P200, upper beta: ∼30–290 ms), progressing through lexical-semantic integration (N400, theta: ∼350–730 ms), and producing behavioral facilitation. Phonological syllable overlap, by contrast, engaged only later oscillatory mechanisms (lower beta: ∼310–590 ms) without modulating early form-sensitive responses, and produced behavioral inhibition. Under parallel activation accounts, both types of syllable overlap should produce overlapping early neural signatures; the selective early sensitivity to orthographic identity, together with the delayed and qualitatively different neural response to phonological overlap, places empirical constraints on the degree of parallelism in orthographic-phonological coordination. While we cannot rule out some degree of cascading activation, the staggered temporal profile is more consistent with the DRC model’s prediction that orthographic analysis precedes phonological assembly (Coltheart et al., 2001) than with the BIAM’s prediction of rapid parallel activation (Grainger & Ziegler, 2011).

Regarding the second research question, the P200 enhancement for orthographic primes reflects facilitated form encoding that precedes lexical access, consistent with the early locus reported for orthographic variables in alphabetic languages (Ferrand & Grainger, 1992, 1993). The absence of early phonological modulation, combined with the later onset of lower beta effects overlapping with the N400 window, suggests that phonological syllable effects may arise at the stage of lexical access. This temporal dissociation provides neural evidence bearing on the processing stage distinction that has been debated primarily on behavioral grounds.

Our findings also reconcile the apparent contradiction in the Korean literature between priming studies showing orthographic facilitation alongside phonological null effects or inhibition (Bae & Yi, 2010; Park, 1996; Tae et al., 2015; Lim et al., 2022) and frequency studies revealing inhibitory phonological neighborhood effects (Kwon et al., 2011; Kwon, 2012).

Critically, the behavioral inhibition observed for phonological primes indicates that phonological codes are actively computed and may engage competitive processes among lexical candidates. This inhibitory pattern converges with the phonological neighborhood effects reported in frequency studies, and is consistent with the sustained lower beta activity in the N400 window, suggesting that both paradigms may tap into overlapping competitive mechanisms at a relatively late processing stage, though the absence of case distinction in Korean means that the early orthographic effects may partly reflect visual identity rather than abstract orthographic processing. Furthermore, these findings bear on the broader phonological recoding debate, in which evidence from multiple languages has supported rapid, obligatory phonological activation (Brysbaert, 2001; Frost, 1998; Lukatela & Turvey, 1994a, 1994b). The present Korean data suggest that the detectability and direction of phonological priming effects are modulated by writing-system properties, as the same phonological overlap that produces facilitation in shallow orthographies produced inhibition here.

We propose that reading systems adapt their processing hierarchy to exploit script-specific regularities. In alpha-syllabic scripts like Korean, the explicit spatial demarcation of syllable boundaries within compact block structures enables a visual-first processing strategy where orthographic parsing provides the primary entry route to the lexicon. This contrasts with alphabetic scripts, where the absence of visual syllable boundaries may necessitate greater reliance on phonological assembly for syllabic parsing. This framework suggests that dual-route models require script-specific parameterization to accommodate the varied processing hierarchies observed across orthographies. The phonological assembly routines formalized in models such as MROM-S (Conrad et al., 2010) and CDP++ (Perry et al., 2010) operate downstream of an initial orthographic parsing stage in Korean, a constraint that future computational modeling efforts should incorporate.

This script-specific processing hierarchy is consistent with broader theoretical proposals that reading systems are shaped by the statistical regularities of their orthography. The psycholinguistic grain size theory (Ziegler & Goswami, 2005) proposes that the functional grain size of sublexical units is determined by the consistency of orthography-to-phonology mappings, with readers of consistent orthographies relying on larger grain sizes. Korean Hangul, which packages graphemes into visually bounded syllable blocks, provides an explicit large-grain orthographic unit that enables direct lexical access without obligatory phonological mediation.

Similarly, Frost (2012) argued that a universal model of reading should accommodate the possibility that phonological computation is not uniformly obligatory across scripts but is modulated by orthographic depth. The present findings are consistent with this proposal that, in Korean’s morphophonemic system, orthographic processing achieves functional priority, while phonological assembly operates as a secondary, downstream process—a configuration that contrasts with the more balanced orthographic-phonological coordination observed in shallow alphabetic orthographies (Frost, 2012; Perfetti & Harris, 2013). This interpretation is also in line with the division-of-labor principle formalized in the triangle model (Harm & Seidenberg, 2004). The model posited that the relative contribution of orthographic and phonological pathways is shaped by the statistical properties of the writing system. In deep orthographies where spelling-to-sound correspondences are inconsistent, the semantic pathway assumes greater functional weight, whereas in systems with reliable orthographic units, the direct orthographic pathway dominates. The compact syllable blocks of Korean orthography provide such reliable orthographic units, supporting efficient direct-route processing consistent with the reduced functional dependence on phonological computation.

### 4.5. Limitations and Future Directions

Several limitations warrant consideration. First, our masked priming design with a brief 50-ms prime duration may have preferentially engaged orthographic processing due to limited time for phonological assembly. Future studies employing longer prime durations or prime awareness manipulations could assess whether phonological effects emerge more robustly under conditions that afford greater processing time, as suggested by the SOA-dependent facilitation observed in French by Ferrand et al. (1996). Notably, Hangul’s orthographic blocks provide immediate syllabic form information, and a brief 50-ms prime may be sufficient to saturate orthographic facilitation while leaving phonological assembly to later stages. Future studies employing systematically varied prime durations (e.g., 50, 100, 150 ms) would help determine the temporal threshold at which phonological priming effects become detectable in Korean. Second, the present study employed visual-visual masked priming exclusively, leaving open the question of whether the observed orthographic facilitation and phonological inhibition pattern generalizes across input modalities. Tae et al. (2017) demonstrated that orthographic priming effects persisted even with auditory primes in a cross-modal paradigm, suggesting that orthographic representations may be activated regardless of input modality. Future studies employing auditory-visual and auditory-auditory priming conditions with concurrent EEG recording could determine whether the temporal dissociation between orthographic and phonological processing observed here reflects a modality-general property of Korean syllable representations or is specific to the visual input channel. Third, our syllable-level analysis leaves open questions about morpheme-level and sentence-level processing, where phonological information may play different functional roles. The inhibitory syllable frequency effects observed in simple nouns (Kwon, 2012) versus facilitatory effects in morphologically complex words (Kim et al., 2023; Kwon et al., 2023) suggest that processing dynamics may differ across lexical structures—a distinction that future ERP and TFR studies with morphologically varied stimuli could address.

## 5. Conclusion

By leveraging the unique orthographic-phonological dissociation afforded by the Korean writing system and employing converging ERP and time-frequency analyses, we observed that orthographic and phonological syllable processing follow distinct spatiotemporal trajectories in Korean visual word recognition. Orthographic syllable overlap facilitates processing through rapid form-based encoding (P200/N/P150 enhancement, upper beta synchronization) followed by efficient lexical-semantic integration (N400 reduction, theta suppression). Phonological syllable overlap, though engaging neural processing at oscillatory levels (sustained lower beta), operates at a later processing stage and produces behavioral inhibition, suggesting lexical competition when orthographic information conflicts. These findings are consistent with a sequential interpretation of orthographic-phonological coordination at the sublexical level, aligning with dual-route cascaded frameworks (Coltheart et al., 2001). Critically, the absence of phonological priming effects at the N250 latency—where the BIAM predicts rapid parallel phonological activation (Grainger & Holcomb, 2009; Grainger & Ziegler, 2011)—places empirical constraints on strong forms of simultaneous parallel activation and obligatory phonological recoding, at least in Korean’s alpha-syllabic writing system. While the N/P150 and N250 component taxonomy established by Grainger et al. (2006) provides a valuable descriptive framework for characterizing the temporal cascade of orthographic and phonological processing, the present data suggest that the functional relationship between these stages is better characterized as sequential rather than parallel, extending this dissociation to a typologically distinct writing system.

## Acknowledgments

This work was supported by the Ministry of Education of the Republic of Korea and the National Research Foundation of Korea (NRF-2023S1A5A2A01074440).

## References

Álvarez, C., Carreiras, M., & Perea, M. (2004). Are syllables phonological units in visual word recognition? Language and Cognitive Processes, 19(3), 427–452. 10.1080/01690960344000242

Andrews, S. (1997). The effect of orthographic similarity on lexical retrieval: Resolving neighborhood conflicts. Psychonomic Bulletin & Review, 4(4), 439–461. 10.3758/BF03214334

Baayen, R. H., Davidson, D. J., & Bates, D. M. (2008). Mixed-effects modeling with crossed random effects for subjects and items. Journal of Memory and Language, 59(4), 390–412. 10.1016/j.jml.2007.12.005

Bae, S., & Yi, K. (2010). Processing of Orthography and Phonology in Korean Word Recognition. Korean Journal of Cognitive and Biological Psychology, 22(3), 369–385. 10.22172/cogbio.2010.22.3.007

Barber, H., Vergara, M., & Carreiras, M. (2004). Syllable-frequency effects in visual word recognition: evidence from ERPs. Neuroreport, 3(15), 545–548. 10.1097/01.wnr.0000111325.38420.80

Bastiaansen, M., Oostenveld, R., Jensen, O., & Hagoort, P. (2008). I see what you mean: Theta power increases are involved in the retrieval of lexical semantic information. Brain and Language, 106(1), 15–28. 10.1016/j.bandl.2007.10.006

Bastiaansen, M., van der Linden, M., ter Keurs, M., Dijkstra, T., & Hagoort, P. (2005). Theta Responses Are Involved in Lexical—Semantic Retrieval during Language Processing. Journal of Cognitive Neuroscience, 17(3), 530–541. 10.1162/0898929053279469

Brysbaert, M. (2001). Prelexical phonological coding of visual words in Dutch: Automatic after all. Memory & Cognition, 29(5), 765–773. 10.3758/BF03200479

Bürkner, P.-C. (2017). brms : An R Package for Bayesian Multilevel Models Using Stan. Journal of Statistical Software, 80, 1. 10.18637/jss.v080.i01

Carreiras, M., Álvarez, C. J. C. J., De Vega, M., Devega, M., De Vega, M., & Devega, M. (1993). Syllable Frequency and Visual Word Recognition in Spanish. Journal of Memory and Language, 32(6), 766–780. 10.1006/jmla.1993.1038

Carreiras, M., & Perea, M. (2002). Masked priming effects with syllabic neighbors in a lexical decision task. Journal of Experimental Psychology: Human Perception and Performance, 28(5), 1228–1242. 10.1037/0096-1523.28.5.1228

Carreiras, M., Perea, M., Vergara, M., & Pollatsek, A. (2009). The time course of orthography and phonology: ERP correlates of masked priming effects in Spanish. Psychophysiology, 46(5), 1113–1122. 10.1111/j.1469-8986.2009.00844.x

Carreiras, M., Vergara, M., & Barber, H. (2005). Early Event-related Potential Effects of Syllabic Processing during Visual Word Recognition. Journal of Cognitive Neuroscience, 17(11), 1803–1817. 10.1162/089892905774589217

Chetail, F., Colin, C., & Content, A. (2012). Electrophysiological markers of syllable frequency during written word recognition in French. Neuropsychologia, 50(14), 3429–3439. 10.1016/j.neuropsychologia.2012.09.044

Choi, W., Lee, C.-H., Kang, J., & Nam, K. (2015). The lexical inhibition of the phonological information in Korean visual word recognition. Korean Journal of Cognitive and Biological Psychology, 27(3), 561–581. 10.22172/cogbio.2015.27.3.011

Coltheart, M., Rastle, K., Perry, C., Langdon, R., & Ziegler, J. (2001). DRC: A dual route cascaded model of visual word recognition and reading aloud. Psychological Review, 108(1), 204–256. 10.1037/0033-295X.108.1.204

Conrad, M., & Jacobs, A. M. (2004). Replicating syllable frequency effects in Spanish in German: One more challenge to computational models of visual word recognition. Language and Cognitive Processes, 19(3), 369–390. 10.1080/01690960344000224

Conrad, M., Tamm, S., Carreiras, M., & Jacobs, A. M. (2010). Simulating syllable frequency effects within an interactive activation framework. In European Journal of Cognitive Psychology (Vol. 22, Number 5, pp. 861–893). Psychology Press Ltd. 10.1080/09541440903356777

Davis, C. J., & Lupker, S. J. (2006). Masked inhibitory priming in English: Evidence for lexical inhibition. Journal of Experimental Psychology: Human Perception and Performance, 32(3), 668–687. 10.1037/0096-1523.32.3.668

Diependaele, K., Ziegler, J. C., & Grainger, J. (2010). Fast phonology and the Bimodal Interactive Activation Model. European Journal of Cognitive Psychology, 22(5), 764–778. 10.1080/09541440902834782

Dominguez, A., De Vega, M., & Cuetos, F. (1997). Lexical Inhibition from Syllabic Units in Spanish Visual Word Recognition. Language and Cognitive Processes, 12(4), 401–422. 10.1080/016909697386790

Engel, A. K., & Fries, P. (2010). Beta-band oscillations — signalling the status quo? Current Opinion in Neurobiology, 20(2), 156–165. 10.1016/j.conb.2010.02.015

Ferrand, L., & Grainger, J. (1992). Phonology and Orthography in Visual Word Recognition: Evidence from Masked Non-Word Priming. The Quarterly Journal of Experimental Psychology Section A, 45(3), 353–372. 10.1080/02724989208250619

Ferrand, L., & Grainger, J. (1993). The time course of orthographic and phonological code activation in the early phases of visual word recognition. Bulletin of the Psychonomic Society, 31(2), 119–122. 10.3758/BF03334157

Frost, R. (2012). Towards a universal model of reading. Behavioral and Brain Sciences, 35(5), 263–279. 10.1017/S0140525X11001841

Goslin, J., Grainger, J., & Holcomb, P. J. (2006). Syllable frequency effects in French visual word recognition: An ERP study. Brain Research, 1115(1), 121–134. 10.1016/j.brainres.2006.07.093

Grainger, J., & Holcomb, P. J. (2009). Watching the Word Go by: On the Time-course of Component Processes in Visual Word Recognition. 10.1111/j.1749-818x.2008.00121.x

Grainger, J., & Jacobs, A. M. (1996). Orthographic Processing in Visual Word Recognition: A Multiple Read-Out Model. Psychological Review, 103(3), 518–565. 10.1037/0033-295X.103.3.518

Grainger, J., Kiyonaga, K., & Holcomb, P. J. (2006). The Time Course of Orthographic and Phonological Code Activation. Psychological Science, 17(12), 1021–1026. 10.1111/j.1467-9280.2006.01821.x

Grainger, J., Muneaux, M., Farioli, F., & Ziegler, J. C. (2005). Effects of Phonological and Orthographic Neighbourhood Density Interact in Visual Word Recognition. The Quarterly Journal of Experimental Psychology Section A, 58(6), 981–998. 10.1080/02724980443000386

Grainger, J., & Ziegler, J. C. (2011). A dual-route approach to orthographic processing. Frontiers in Psychology, 2(APR), 1–13. 10.3389/fpsyg.2011.00054

Harm, M. W., & Seidenberg, M. S. (2004). Computing the Meanings of Words in Reading: Cooperative Division of Labor Between Visual and Phonological Processes. Psychological Review, 111(3), 662–720. 10.1037/0033-295X.111.3.662

Holcomb, P. J., Akers, E. M., Midgley, K. J., & Emmorey, K. (2024). Orthographic and Phonological Code Activation in Deaf and Hearing Readers. Journal of Cognition, 7(1). 10.5334/joc.326

Holcomb, P. J., & Grainger, J. (2006). On the Time Course of Visual Word Recognition: An Event-related Potential Investigation using Masked Repetition Priming. Journal of Cognitive Neuroscience, 18(10), 1631–1643. 10.1162/jocn.2006.18.10.1631

Hutzler, F., Bergmann, J., Conrad, M., Kronbichler, M., Stenneken, P., & Jacobs, A. M. (2004). Inhibitory effects of first syllable-frequency in lexical decision: an event-related potential study. Neuroscience Letters, 372(3), 179–184. 10.1016/j.neulet.2004.07.050

Jin, R., Lee, H., & Choi, W. (2018). Are they real neighbors?: Null effects of syllabic neighbors in Korean word recognition. Korean Journal of Cognitive and Biological Psychology, 30(3), 211–223. 10.22172/cogbio.2018.30.3.001

Kilavik, B. E., Zaepffel, M., Brovelli, A., MacKay, W. A., & Riehle, A. (2013). The ups and downs of beta oscillations in sensorimotor cortex. Experimental Neurology, 245(1), 15–26. 10.1016/j.expneurol.2012.09.014

Kim, J., Lee, S., Kim, S., & Nam, K. (2023). Syllable frequency effect in visual word recognition: a regression study on morphologically simple and complex Korean words. Korean Journal of Cognitive and Biological Psychology, 35(4), 303–335. 10.22172/cogbio.2023.35.4.004

Klimesch, W. (1999). EEG alpha and theta oscillations reflect cognitive and memory performance: a review and analysis. Brain Research Reviews, 29(2–3), 169–195. 10.1016/S0165-0173(98)00056-3

Klimesch, W., Doppelmayr, M., Wimmer, H., Gruber, W., Röhm, D., Schwaiger, J., & Hutzler, F. (2001). Alpha and beta band power changes in normal and dyslexic children. Clinical Neurophysiology, 112(7), 1186–1195. 10.1016/S1388-2457(01)00543-0

Kwon, S., Kim, J., Lee, S., & Nam, K. (2023). The Facilitative Effect of First Syllable Frequency during Visual Recognition of Korean Noun Eojeols. The Korean Journal of Cognitive and Biological Psychology, 35(2), 93–106. 10.22172/cogbio.2023.35.2.004

Kwon, S., Lee, S., Kim, J., & Nam, K. (2024). The time course of syllable frequency effects in the visual recognition of Korean morphologically complex nouns: an ERP study. Frontiers in Language Sciences, 3, 1477606. 10.3389/flang.2024.1477606

Kwon, Y. (2012). The Dissociation of Syllabic Token and Type Frequency Effect in Lexical Decision Task. Korean Journal of Cognitive and Biological Psychology, 24(4), 315–333. 10.22172/cogbio.2012.24.4.002

Kwon, Y., & Lee, Y. (2015). The Source of the Syllable Frequency Effect During Visual Word Recognition : Event-Related Brain Potential Stud. Journal of Language Sciences, 22(4), 1–17. 10.14384/kals.2015.22.4.001

Kwon, Y., & Lee, Y. (2017). The Reason for the Absence of the Syllable Frequency Effect in Korean: Behavioral and ERP Evidences from Morphological Syllable. Journal of The Korean Data Analysis Society, 19(1), 465–476.

Kwon, Y., Lee, Y., & Nam, K. (2011). The different P200 effects of phonological and orthographic syllable frequency in visual word recognition in Korean. Neuroscience Letters, 501(2), 117–121. 10.1016/j.neulet.2011.06.060

Kwon, Y., Nam, K., & Lee, Y. (2012). ERP index of the morphological family size effect during word recognition. Neuropsychologia, 50(14), 3385–3391. 10.1016/j.neuropsychologia.2012.09.041

Lee, C. H., & Taft, M. (2009). Are onsets and codas important in processing letter position? A comparison of TL effects in English and Korean. Journal of Memory and Language, 60(4), 530–542. 10.1016/j.jml.2009.01.002

Lee, M. D., & Wagenmakers, E.-J. (2012). Bayesian Cognitive Modeling: A Practical Course. Cambridge University Press.

Lee, S., Kim, S., Kim, J., Kwon, S., Lee, E.-H., & Nam, K. (2023). The Facilitative Effect of the First Syllable Token Frequency in Visual Recognition of Korean Predicate Eojeols. The Korean Journal of Cognitive and Biological Psychology, 2023(4), 337–345. 10.22172/cogbio.2023.35.4.005

Lee, Y., Kwon, Y., & Lee, C. (2016). The Event Related Potential Evidence for the Orthographic and Phonological Priming in Korean Visual Word Recognition. Journal of The Korean Data Analysis Society, 18(4), 2093–2105.

Lewis, A. G., & Bastiaansen, M. (2015). A predictive coding framework for rapid neural dynamics during sentence-level language comprehension. Cortex, 68, 155–168. 10.1016/j.cortex.2015.02.014

Lewis, A. G., Wang, L., & Bastiaansen, M. (2015). Fast oscillatory dynamics during language comprehension: Unification versus maintenance and prediction? Brain and Language, 148, 51–63. 10.1016/j.bandl.2015.01.003

Lim, C., Baek, H., Hoon Kim, T., & Choi, W. (2022). Activation of Phonological and Orthographic information during Korean Visual Word Recognition: Evidence from a Meta-analysis and a Priming Study. The Korean Journal of Cognitive and Biological Psychology, 34(4), 221–236. 10.22172/cogbio.2022.34.4.002

Luce, P. A., & Pisoni, D. B. (1998). Recognizing Spoken Words: The Neighborhood Activation Model. Ear Hear, 19(1), 1–36.

Lukatela, G., Eaton, T., Lee, C., & Turvey, M. T. (2001). Does visual word identification involve a sub-phonemic level? Cognition, 78(3), B41–B52. 10.1016/S0010-0277(00)00121-9

Lukatela, G., Frost, S. J., & Turvey, M. T. (1998). Phonological Priming by Masked Nonword Primes in the Lexical Decision Task. Journal of Memory and Language, 39(4), 666–683. 10.1006/jmla.1998.2599

Lukatela, G., & Turvey, M. T. (1994a). Visual lexical access is initially phonological: 2. Evidence from phonological priming by homophones and pseudohomophones. Journal of Experimental Psychology: General, 123(4), 331–353. 10.1037/0096-3445.123.4.331

Lukatela, G., & Turvey, M. T. (1994b). Visual lexical access is initially phonological: I. Evidence from associative priming by words, homophones, and pseudohomophones. Journal of Experimental Psychology: General, 123(2), 107–128. 10.1037/0096-3445.123.2.107

Lupker, S. J., & Colombo, L. (1994). Inhibitory effects in form priming: Evaluating a phonological competition explanation. Journal of Experimental Psychology: Human Perception and Performance, 20(2), 437–451. 10.1037/0096-1523.20.2.437

Mai, G., Minett, J. W., & Wang, W. S. Y. (2016). Delta, theta, beta, and gamma brain oscillations index levels of auditory sentence processing. NeuroImage, 133(7), 516–528. 10.1016/j.neuroimage.2016.02.064

Mathey, S., & Zagar, D. (2002). Similarity in Visual Word Recognition: The Effect of Syllabic Neighborood in French. *Current Psychology Letters: Behaviour*, Brain & Cognition, (8), 107–121. 10.4000/cpl.210

McClelland, J. L., & Rumelhart, D. E. (1981). An interactive activation model of context effects in letter perception: I. An account of basic findings. Psychological Review, 88(5), 375–407. 10.1037/0033-295X.88.5.375

Oostenveld, R., Fries, P., Maris, E., & Schoffelen, J. M. (2011). FieldTrip: Open source software for advanced analysis of MEG, EEG, and invasive electrophysiological data. Computational Intelligence and Neuroscience, 2011. 10.1155/2011/156869

Pae, H. K. (2011). Is Korean a syllabic alphabet or an alphabetic syllabary. Writing Systems Research, 3(2), 103–115. 10.1093/wsr/wsr002

Pae, H. K., Bae, S., & Yi, K. (2020). Lexical properties influencing visual word recognition in Hangul. Reading and Writing, 33(9), 2391–2412. 10.1007/s11145-020-10042-4

Park, K. (1996). The Role of Phonology in Hangul Word Recognition. The Korean Journal of Experimental and Cognitive Psychology, 8(1), 25–44.

Perea, M., & Carreiras, M. (1998). Effects of Syllable Frequency and Syllable Neighborhood Frequency in Visual Word Recognition. Journal of Experimental Psychology: Human Perception and Performance, 24(1), 134–144. 10.1037/0096-1523.24.1.134

Perfetti, C. A., & Harris, L. N. (2013). Universal Reading Processes Are Modulated by Language and Writing System. Language Learning and Development, 9(4), 296–316. 10.1080/15475441.2013.813828

Perry, C., Ziegler, J. C., & Zorzi, M. (2007). Nested incremental modeling in the development of computational theories: The CDP+ model of reading aloud. Psychological Review, 114(2), 273–315. 10.1037/0033-295X.114.2.273

Perry, C., Ziegler, J. C., & Zorzi, M. (2010). Beyond single syllables: Large-scale modeling of reading aloud with the Connectionist Dual Process (CDP++) model. Cognitive Psychology, 61(2), 106–151. 10.1016/j.cogpsych.2010.04.001

Pfurtscheller, G., & Lopes da Silva, F. H. (1999). Event-related EEG/MEG synchronization and desynchronization: basic principles. Clinical Neurophysiology, 110(11), 1842–1857. 10.1016/S1388-2457(99)00141-8

Piai, V., Roelofs, A., & Maris, E. (2014). Oscillatory brain responses in spoken word production reflect lexical frequency and sentential constraint. Neuropsychologia, 53(1), 146–156. 10.1016/j.neuropsychologia.2013.11.014

R Core Team. (2025). R: A Language and Environment for Statistical Computing. https://www.R-project.org/

Rastle, K., & Brysbaert, M. (2006). Masked phonological priming effects in English: Are they real? Do they matter? Cognitive Psychology, 53(2), 97–145. 10.1016/j.cogpsych.2006.01.002

Simpson, G. B., & Kang, H. (2004). Syllable processing in alphabetic Korean. Reading and Writing, 17(1–2), 137–151. 10.1023/b:read.0000013808.65933.a1

Spitzer, B., & Haegens, S. (2017). Beyond the Status Quo: A Role for Beta Oscillations in Endogenous Content (Re)Activation. Eneuro, 4(4), ENEURO.0170-17.2017. 10.1523/ENEURO.0170-17.2017

Tae, J., Lee, C., & Lee, Y. (2015). The Effect of the Orthographic and Phonological Priming in Korean Visual Word Recognition. Korean Journal of Cognitive Science, 26(1), 1–26.

Tae, J., Lee, Y., & Kwon, Y. (2017). The Role of Phonological Information on Visual Word Recognition through Auditory-visual Cross Modal Priming. Journal of Language Sciences, 24(1), 175–189. 10.14384/kals.2017.24.1.175

Wagenmakers, E. J., Lodewyckx, T., Kuriyal, H., & Grasman, R. (2010). Bayesian hypothesis testing for psychologists: A tutorial on the Savage–Dickey method. Cognitive Psychology, 60(3), 158–189. 10.1016/j.cogpsych.2009.12.001

Wang, L., Jensen, O., van den Brink, D., Weder, N., Schoffelen, J.-M. M., Magyari, L., Hagoort, P., & Bastiaansen, M. (2012). Beta oscillations relate to the N400m during language comprehension. Human Brain Mapping, 33(12), 2898–2912. 10.1002/hbm.21410

Weiss, S., & Mueller, H. M. (2012). “Too Many betas do not Spoil the Broth”: The Role of Beta Brain Oscillations in Language Processing. Frontiers in Psychology, 3(JUN). 10.3389/fpsyg.2012.00201

Ziegler, J. C., & Goswami, U. (2005). Reading Acquisition, Developmental Dyslexia, and Skilled Reading Across Languages: A Psycholinguistic Grain Size Theory. Psychological Bulletin, 131(1), 3–29. 10.1037/0033-2909.131.1.3

