## Supplementary Materials for "Rapid Orthographic and Delayed Phonological Processing: ERP and Oscillatory Evidence from Masked Priming in Korean"

### S1. P200 Full Bayesian LMM (Both ROIs Combined)

A Bayesian LMM was fitted to P200 amplitudes (150–250 ms) with ROI (FrontoCentral, CentroParietal), Condition, and their interaction as fixed factors; List was included as a covariate. ROI was sum-coded (±0.5); Condition was reference-coded with Unrelated as the baseline. The ROI main effect was large and negative (β = −0.706 μV), reflecting higher P200 amplitude at FrontoCentral sites. No ROI × Condition interaction reached moderate evidence (Evid.Ratio₀ ≥ 5 for all interaction terms).

**Table S1**

*P200 Full-Model Fixed Effects (Both ROIs Combined)*

| **Parameter** | **Contrast** | **β (SE)** | **95% CI** | **R̂** | **pd** | **Evid.Ratio₁** | **Evid.Ratio₀** |
| --- | --- | --- | --- | --- | --- | --- | --- |
| Condition₁ | O vs. UR | 0.233  (0.102) | [0.033, 0.432] | 1.00 | 0.989 | 90.9 | 0.82 |
| Condition₂ | P vs. UR | 0.099  (0.100) | [−0.099, 0.296] | 1.00 | 0.841 | 5.29 | 6.56 |
| Condition₁₋₂ | O vs. P | 0.134  (0.102) | [−0.064, 0.335] | 1.00 | 0.907 | 9.74 | 5.79 |
| ROI × Cond₁ | ROI ×  (O vs. UR) | −0.112 (0.099) | [−0.308, 0.079] | 1.00 | 0.867 | 6.52 | 5.43 |
| ROI × Cond₂ | ROI ×  (P vs. UR) | −0.053 (0.098) | [−0.246, 0.141] | 1.00 | 0.703 | 2.37 | 8.77 |
| ***Random Effects*** | | | | | | | |
| Subject |  | SD = 0.788 | — | — | — | — | — |
| Item |  | SD = 0.390 | — | — | — | — | — |

*Note.* Condition₁ = Orthographic vs. Unrelated (O vs. UR); Condition₂ = Phonological vs. Unrelated (P vs. UR); Condition₁₋₂ = Orthographic vs. Phonological (O vs. P). β = posterior mean (μV); SE = posterior SD; 95% CI = 95% credible interval; pd = probability of the dominant direction. Evid.Ratio₁ = pd/(1−pd): directional evidence ratio (prior-insensitive). Evid.Ratio₀ = Savage–Dickey BF₀₁: evidence for the point null H₀: β = 0 (prior-sensitive); values < 1 indicate evidence for the non-zero alternative (BF₁₀ = 1/Evid.Ratio₀). SD = posterior standard deviation of random intercept.

### S2. P200 ROI-Specific Bayesian LMM — FrontoCentral

A Bayesian LMM was fitted to P200 amplitudes at FrontoCentral sites (Fz, F1, F2, FC1, FC2, Cz), the theoretically predicted ROI for early orthographic processing. Orthographic primes produced strongly larger P200 amplitudes than unrelated primes (Evid.Ratio₁ = 139.4, Evid.Ratio₀ = 0.40, indicating converging evidence for a positive non-zero effect). No reliable phonological priming effect was observed (Evid.Ratio₁ = 6.24, Evid.Ratio₀ = 3.96).

**Table S2**

*P200 FrontoCentral ROI-Specific Fixed Effects (Primary)*

| **Parameter** | **Contrast** | **β (SE)** | **95% CI** | **R̂** | **pd** | **Evid.Ratio₁** | **Evid.Ratio₀** |
| --- | --- | --- | --- | --- | --- | --- | --- |
| Condition₁ | O vs. UR | 0.233  (0.102) | [0.069, 0.622] | 1.00 | 0.993 | 139.4 | 0.40 |
| Condition₂ | P vs. UR | 0.099  (0.100) | [−0.124, 0.420] | 1.00 | 0.862 | 6.24 | 3.96 |
| Condition₁₋₂ | O vs. P | 0.134  (0.102) | [−0.078, 0.470] | 1.00 | 0.914 | 10.63 | 3.92 |
| ***Random Effects*** | | | | | | | |
| Subject |  | SD = 1.376 | — | — | — | — | — |
| Item |  | SD = 0.888 | — | — | — | — | — |

*Note.* Condition₁ = Orthographic vs. Unrelated (O vs. UR); Condition₂ = Phonological vs. Unrelated (P vs. UR); Condition₁₋₂ = Orthographic vs. Phonological (O vs. P). β = posterior mean (μV); SE = posterior SD; 95% CI = 95% credible interval; pd = probability of the dominant direction. Evid.Ratio₁ = pd/(1−pd): directional evidence ratio (prior-insensitive). Evid.Ratio₀ = Savage–Dickey BF₀₁: evidence for the point null H₀: β = 0 (prior-sensitive); values < 1 indicate evidence for the non-zero alternative (BF₁₀ = 1/Evid.Ratio₀). SD = posterior standard deviation of random intercept.

### S3. P200 ROI-Specific Bayesian LMM — CentroParietal

A Bayesian LMM was fitted to P200 amplitudes at CentroParietal sites (CPz, CP1, CP2, P1, P2, Pz). This ROI was not the theoretically predicted site for P200 modulation. All 95% CIs included zero and Evid.Ratio₀ values indicated moderate-to-strong evidence for the null hypothesis (Evid.Ratio₀ = 5.03–10.69).

**Table S3**

*P200 CentroParietal ROI-Specific Fixed Effects (Non-Hypothesized)*

| **Parameter** | **Contrast** | **β (SE)** | **95% CI** | **R̂** | **pd** | **Evid.Ratio₁** | **Evid.Ratio₀** |
| --- | --- | --- | --- | --- | --- | --- | --- |
| Condition₁ | O vs. UR | 0.118  (0.128) | [−0.135, 0.372] | 1.00 | 0.826 | 4.75 | 5.03 |
| Condition₂ | P vs. UR | 0.081  (0.130) | [−0.170, 0.335] | 1.00 | 0.734 | 2.76 | 6.38 |
| Condition₁₋₂ | O vs. P | 0.038  (0.132) | [−0.225, 0.292] | 1.00 | 0.619 | 1.63 | 10.69 |
| ***Random Effects*** | | | | | | | |
| Subject |  | SD = 1.376 | — | — | — | — | — |
| Item |  | SD = 0.888 | — | — | — | — | — |

*Note.* Condition₁ = Orthographic vs. Unrelated (O vs. UR); Condition₂ = Phonological vs. Unrelated (P vs. UR); Condition₁₋₂ = Orthographic vs. Phonological (O vs. P). β = posterior mean (μV); SE = posterior SD; 95% CI = 95% credible interval; pd = probability of the dominant direction. Evid.Ratio₁ = pd/(1−pd): directional evidence ratio (prior-insensitive). Evid.Ratio₀ = Savage–Dickey BF₀₁: evidence for the point null H₀: β = 0 (prior-sensitive); values < 1 indicate evidence for the non-zero alternative (BF₁₀ = 1/Evid.Ratio₀). SD = posterior standard deviation of random intercept.

### S4. N400 Full Bayesian LMM (Both ROIs Combined)

A Bayesian LMM was fitted to N400 amplitudes (350–550 ms) with the same structure as the P200 full model. The ROI main effect was large and positive (β = 1.174 μV), reflecting greater N400 amplitude at CentroParietal sites. The orthographic condition effect showed high directional probability (pd = 0.961, Evid.Ratio₁ = 24.6) but also yielded only anecdotal evidence against the point null (Evid.Ratio₀ = 2.08), reflecting a small-to-moderate effect consistent in direction yet uncertain in magnitude. The ROI × Condition₁ interaction similarly showed moderate directional probability (Evid.Ratio₁ = 13.10, Evid.Ratio₀ = 3.63), motivating ROI-specific analyses.

**Table S4**

*N400 Full-Model Fixed Effects (Both ROIs Combined)*

| **Parameter** | **Contrast** | **β (SE)** | **95% CI** | **R̂** | **pd** | **Evid.Ratio₁** | **Evid.Ratio₀** |
| --- | --- | --- | --- | --- | --- | --- | --- |
| Condition₁ | O vs. UR | 0.175  (0.099) | [−0.021, 0.368] | 1.00 | 0.961 | 24.6 | 2.08 |
| Condition₂ | P vs. UR | 0.043  (0.099) | [−0.148, 0.238] | 1.00 | 0.666 | 2.00 | 9.23 |
| Condition₁₋₂ | O vs. P | 0.133  (0.101) | [−0.066, 0.331] | 1.00 | 0.905 | 9.50 | 5.77 |
| ROI × Cond₁ | ROI ×  (O vs. UR) | 0.140  (0.096) | [−0.048, 0.331] | 1.00 | 0.929 | 13.10 | 3.63 |
| ROI × Cond₂ | ROI ×  (P vs. UR) | −0.044 (0.097) | [−0.235, 0.143] | 1.00 | 0.671 | 2.04 | 9.01 |
| ***Random Effects*** | | | | | | | |
| Subject |  | SD = 0.812 | — | — | — | — | — |
| Item |  | SD = 0.529 | — | — | — | — | — |

*Note.* Condition₁ = Orthographic vs. Unrelated (O vs. UR); Condition₂ = Phonological vs. Unrelated (P vs. UR); Condition₁₋₂ = Orthographic vs. Phonological (O vs. P). β = posterior mean (μV); SE = posterior SD; 95% CI = 95% credible interval; pd = probability of the dominant direction. Evid.Ratio₁ = pd/(1−pd): directional evidence ratio (prior-insensitive). Evid.Ratio₀ = Savage–Dickey BF₀₁: evidence for the point null H₀: β = 0 (prior-sensitive); values < 1 indicate evidence for the non-zero alternative (BF₁₀ = 1/Evid.Ratio₀). SD = posterior standard deviation of random intercept.

### S5. N400 ROI-Specific Bayesian LMM — CentroParietal

A Bayesian LMM was fitted to N400 amplitudes at CentroParietal sites (CPz, CP1, CP2, P1, P2, Pz), the theoretically predicted ROI for lexical-semantic N400 effects. Orthographic primes produced a robust N400 reduction compared to both unrelated primes (Evid.Ratio₁ = 116.6, Evid.Ratio₀ = 0.45) and phonological primes (Evid.Ratio₁ = 81.0, Evid.Ratio₀ = 0.86). Phonological primes did not differ from unrelated primes (Evid.Ratio₁ = 1.30, Evid.Ratio₀ = 7.73).

**Table S5**

*N400 CentroParietal ROI-Specific Fixed Effects (Primary)*

| **Parameter** | **Contrast** | **β (SE)** | **95% CI** | **R̂** | **pd** | **Evid.Ratio₁** | **Evid.Ratio₀** |
| --- | --- | --- | --- | --- | --- | --- | --- |
| Condition₁ | O vs. UR | 0.317 (0.132) | [0.059, 0.576] | 1.00 | 0.992 | 116.6 | 0.45 |
| Condition₂ | P vs. UR | 0.019 (0.132) | [−0.238, 0.277] | 1.00 | 0.566 | 1.30 | 7.73 |
| Condition₁₋₂ | O vs. P | 0.298 (0.132) | [0.040, 0.558] | 1.00 | 0.988 | 81.0 | 0.86 |
| ***Random Effects*** | | | | | | | |
| Subject |  | SD = 1.397 | — | — | — | — | — |
| Item |  | SD = 1.096 | — | — | — | — | — |

*Note.* Condition₁ = Orthographic vs. Unrelated (O vs. UR); Condition₂ = Phonological vs. Unrelated (P vs. UR); Condition₁₋₂ = Orthographic vs. Phonological (O vs. P). β = posterior mean (μV); SE = posterior SD; 95% CI = 95% credible interval; pd = probability of the dominant direction. Evid.Ratio₁ = pd/(1−pd): directional evidence ratio (prior-insensitive). Evid.Ratio₀ = Savage–Dickey BF₀₁: evidence for the point null H₀: β = 0 (prior-sensitive); values < 1 indicate evidence for the non-zero alternative (BF₁₀ = 1/Evid.Ratio₀). SD = posterior standard deviation of random intercept.

### S6. N400 ROI-Specific Bayesian LMM — FrontoCentral

A Bayesian LMM was fitted to N400 amplitudes at FrontoCentral sites (Fz, F1, F2, FC1, FC2, Cz). No reliable condition effects were observed: all 95% CIs included zero, and Evid.Ratio₀ values indicated moderate-to-strong evidence for the null (Evid.Ratio₀ = 6.30–10.30), consistent with the canonical centro-parietal topography of the N400.

**Table S6**

*N400 FrontoCentral ROI-Specific Fixed Effects (Non-Hypothesized)*

| **Parameter** | **Contrast** | **β (SE)** | **95% CI** | **R̂** | **pd** | **Evid.Ratio₁** | **Evid.Ratio₀** |
| --- | --- | --- | --- | --- | --- | --- | --- |
| Condition₁ | O vs. UR | 0.044  (0.127) | [−0.206, 0.289] | 1.00 | 0.637 | 1.75 | 7.44 |
| Condition₂ | P vs. UR | 0.077  (0.128) | [−0.184, 0.316] | 1.00 | 0.728 | 2.68 | 6.30 |
| Condition₁₋₂ | O vs. P | −0.033 (0.129) | [−0.285, 0.222] | 1.00 | 0.607 | 1.55 | 10.30 |
| ***Random Effects*** | | | | | | | |
| Subject |  | SD = 1.397 | — | — | — | — | — |
| Item |  | SD = 1.096 | — | — | — | — | — |

*Note.* Condition₁ = Orthographic vs. Unrelated (O vs. UR); Condition₂ = Phonological vs. Unrelated (P vs. UR); Condition₁₋₂ = Orthographic vs. Phonological (O vs. P). β = posterior mean (μV); SE = posterior SD; 95% CI = 95% credible interval; pd = probability of the dominant direction (always ≥ 0.5). Evid.Ratio₁ = pd/(1−pd): directional evidence ratio (prior-insensitive). Evid.Ratio₀ = Savage–Dickey BF₀₁: evidence for the point null H₀: β = 0 (prior-sensitive); values < 1 indicate evidence for the non-zero alternative (BF₁₀ = 1/Evid.Ratio₀). R̂ ≤ 1.01 confirmed for all parameters. SD = posterior standard deviation of random intercept.
